# Expression dynamics of long non-coding RNAs during imaginal disc development and regeneration in *Drosophila*

**DOI:** 10.1101/2024.03.19.585729

**Authors:** Carlos Camilleri-Robles, Raziel Amador, Marcel Tiebe, Aurelio A Teleman, Florenci Serras, Roderic Guigó, Montserrat Corominas

## Abstract

The discovery of functional long non-coding RNAs (lncRNAs) changed their initial concept as transcriptional noise. LncRNAs have been found to participate in the regulation of multiple biological processes, including chromatin structure, gene expression, splicing, and mRNA degradation and translation. However, functional studies of lncRNAs are hindered by the usual lack of phenotypes upon deletion or inhibition. Here, we used *Drosophila* imaginal discs as a model system to identify lncRNAs involved in development and regeneration. We examined a subset of lncRNAs expressed in the wing, leg, and eye disc development. Additionally, we analyzed transcriptomic data from regenerating wing discs to profile the expression pattern of lncRNAs during tissue repair. We focused on the lncRNA *CR40469*, which is upregulated during regeneration. We generated *CR40469* mutant flies that developed normally but showed impaired wing regeneration upon the induction of cell death. The ability of these mutants to regenerate was restored by the ectopic expression of *CR40469*. Furthermore, we found that the lncRNA *CR34335* has a high degree of sequence similarity with *CR40469* and can partially compensate for its function during regeneration in the absence of *CR40469*. Our findings point to a potential role of the lncRNA *CR40469* in *trans* during the response to damage in the wing imaginal disc.

## Introduction

Long non-coding RNAs (lncRNAs) are defined as transcripts longer than 200 nucleotides that lack protein-coding potential. They have been mostly identified in high-throughput transcriptomic studies and are quite abundant in metazoa (Derrien et al. 2012; Pauli et al. 2012; Brown et al. 2014). They tend to show low sequence conservation (Derrien et al. 2012; Brown et al. 2014), although they may show other conservation signatures such as positional conservation or synteny, structural conservation, or functional convergence (Ramírez-Colmenero et al. 2020). Many do not produce observable phenotypes upon mutation (Liu et al. 2017; Ramilowski et al. 2020), and the vast majority have not been assigned to a putative function (Lee et al. 2019). Moreover, the expression of lncRNAs is generally lower and highly tissue- and stage-specific when compared to that of protein-coding genes (Derrien et al. 2012; Brown et al. 2014; Washietl et al. 2014), hampering their proper characterization. In recent years, a small proportion of annotated lncRNAs have been functionally characterized and described to participate in the regulation of almost every step of gene expression (Statello et al. 2021). Multiple lncRNAs are known to regulate the expression of overlapping or nearby protein-coding genes (Engreitz et al. 2016; Pérez-Lluch et al. 2020), but some are also capable of influencing the expression of genomically distant genes in *trans* (Lee et al. 2016; Tichon et al. 2016). LncRNAs can interact with DNA, RNAs, proteins, and even membrane lipids, showing diverse subcellular distributions (Wilk et al. 2016; Krause 2018).

LncRNAs are involved in multiple developmental processes, including the regulation of Hox genes (Rinn et al. 2007), the regulation of chromatin accessibility in dosage compensation mechanisms (Loda and Heard 2019), and the development of organs such as the brain (Bernard et al. 2010) and heart (Klattenhoff et al. 2013). Alterations in the expression of lncRNAs have also been described for several diseases, including cancer (Huarte 2015; Carlevaro-Fita et al. 2020), cardiovascular diseases (Poller et al. 2018) and neurological disorders (Sunwoo et al. 2017). In fact, the public database LncRNADisease v2.0 currently describes more than 1,700 experimentally validated lncRNA-disease associations (Bao et al. 2019), highlighting lncRNAs as potential tools to understand the mechanisms and prognosis of multiple diseases. Moreover, changes in lncRNA expression have been found in the hypoxia response pathway and in response to various stress conditions including oxidative stress, heat-shock and DNA damage (Valadkhan and Valencia-Hipólito 2015).

The number of annotated lncRNAs varies greatly from one species to another. While the numbers of annotated protein-coding genes and lncRNAs are similar in humans and mice, the number of lncRNAs is considerably lower in *Drosophila melanogaster* and *Caenorhabditis elegans* (Camilleri-Robles et al. 2022). Despite this, some lncRNA features are shared between mammals and *Drosophila*, including the transcript length and the proportion of genic/intergenic lncRNAs (Camilleri-Robles et al. 2022). Moreover, some lncRNAs function similarly in different species, such as the lncRNAs *Xist* and *roX1/roX2*, which recruit the chromatin-modifying complexes that drive the dosage compensation mechanisms in mammals and flies, respectively (Samata and Akhtar 2018; Loda and Heard 2019).

The expression of lncRNAs has been extensively characterized throughout *Drosophila* embryogenesis and larval development, revealing a highly temporally restricted expression profile, especially at the late embryonic and late larval stages, which are critical times for developmental transitions (Chen et al. 2016; Schor et al. 2018). The subcellular localization of 103 lncRNAs has also been determined over the course of embryogenesis and in the late third instar larval tissues (Wilk et al. 2016). However, there is no detailed information regarding lncRNA expression in imaginal discs at different stages of development or after damage.

*Drosophila* imaginal discs are larval epithelial sacs that give rise to the adult structures after differentiation during metamorphosis. Each imaginal disc develops from a cluster of a few cells in the embryo and its morphology and size matures during the larval stages. Mature discs undergo major morphogenetic events during metamorphosis (Beira and Paro 2016). Third instar larval imaginal discs also present a high capacity to regenerate following a physical injury (Bryant 1971; Schubiger 1971) or after genetic ablation (Smith-Bolton et al. 2009; Bergantiños et al. 2010; Hariharan and Serras 2017). Upon damage, early signals that include calcium waves and reactive oxygen species (ROS) are propagated from the dying cells to the neighboring living cells, activating a series of signaling pathways that are required for wound healing and regenerative growth (Santabárbara-Ruiz et al. 2015; Fogarty et al. 2016; Esteban-Collado et al. 2021). The activation of these pathways leads to a burst of active transcription at the early stages of regeneration (Vizcaya-Molina et al. 2018), activating, among others, the transcription factor *Ets21C*, which is described to orchestrate a regeneration-specific gene regulatory network (Worley et al. 2023).

In this work, we analyzed the expression of lncRNAs in different imaginal discs during larval and pupal stages, as well as in the regeneration of wing discs. We found that the intergenic lncRNA *CR40469* is required for regeneration, but not in development. The recovery of the regeneration capacity of *CR40469* mutants with the addition of ectopic *CR40469* led us to hypothesize about a putative *trans*-acting role after damage. Additionally, we found that *CR40469* and a second lncRNA, *CR34335*, share a high sequence similarity. Despite showing opposite expression profiles, the ectopic expression of *CR34335* also rescued the regeneration capacity of *CR40469* mutants, suggesting that it may compensate for the function of *CR40469* in its absence.

## Results

### Expression of lncRNAs in developing imaginal discs

We analyzed the expression profiles of the 2,455 lncRNAs annotated in the *Drosophila* genome (FlyBase genome annotation version r6.29) in imaginal discs and compared them with those of the protein-coding genes (PCGs) using available transcriptomic data from wing, leg and eye discs at three different developmental stages (Ruiz-Romero et al. 2022): third instar larvae (L3), early pupae (EP) and late pupae (LP) (Fig. 1A). The numbers of expressed lncRNAs and PCGs were similar across all the tissues and stages: ∼200 expressed lncRNAs and ∼8,000 expressed PCGs per condition, which represent ∼8% of annotated lncRNAs and 57% of annotated PCGs, respectively (Fig. 1B). In accordance with previous observations, the expression levels of lncRNAs were remarkably lower compared to those of PCGs (Supplemental Fig. S1A-A’’). No differences were observed in terms of transcript length, number of exons or GC content comparing expressed and non-expressed lncRNAs (Supplemental Fig. S1B-E). Most expressed lncRNAs and PCGs were present in the wing, leg and eye discs at the three developmental stages studied, highlighting the genetic similarity of these tissues (Fig. 1C,D). However, we observed a higher tissue- (Fig. 1C) and stage-specificity (Fig. 1D) for lncRNAs compared to PCGs. The stage-specificity was higher at the LP stage for lncRNAs and PCGs, consistent with the morphogenetic events taking place during metamorphosis (Chen et al. 2016). To get insight into the expression profiles of lncRNAs and PCGs during imaginal disc development, we standardized the expression values of each gene across all developmental samples. For PCGs, we identified 4 major gene clusters: (1) genes more expressed at the LP stage, (2) genes more expressed at the L3 and EP stages of wing and leg discs, (3) genes more expressed in the eye, particularly at the LP stage, and (4) genes more expressed at the L3 and EP stages of eye discs (Fig. 1E). On the contrary, lncRNAs formed 2 main clusters: (1) lncRNAs more expressed at LP stages, and (2) lncRNAs more expressed at L3 and EP stages (Fig. 1F).

**Figure 1.**
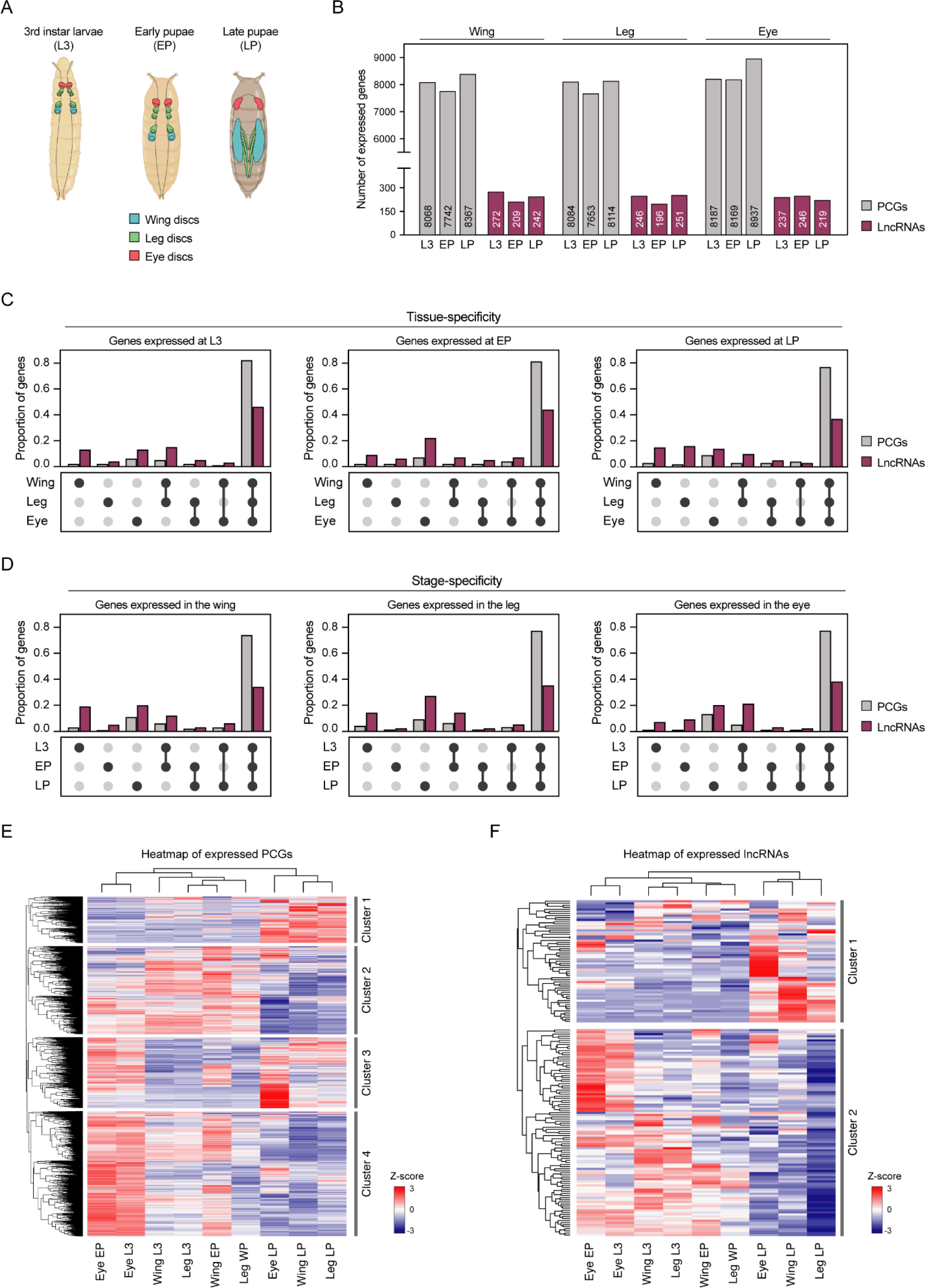
Expressed lncRNAs and PCGs in *Drosophila* imaginal discs. (A) Representation of the developmental samples used for the gene expression analysis. (B) Number of protein-coding genes (PCGs) and lncRNAs expressed in the wing, leg and eye discs in third instar larvae (L3), early pupae (EP) and late pupae (LP). (C) Tissue specificity of PCGs and lncRNAs in L3 (left), EP (middle) and LP (right). (D) Stage specificity of PCGs and lncRNAs in the wing (left), leg (middle) and eye discs (right). (E-F) Expression of (E) PCGs or (F) lncRNAs in development. Gene expression is normalized to Z-score values. Genes not expressed in any sample were not considered. Complete hierarchical clustering was used to identify the gene clusters.

We next classified the expressed lncRNAs as genic (if they were in the exonic or intronic regions of PCGs) or intergenic, and paired each lncRNA with their closest PCG neighbor, independently of the direction of transcription. In this way, intergenic lncRNAs were paired with their closest PCG, while genic lncRNAs were associated with their overlapping PCG (Fig. 2A). Due to the high compactness of the *Drosophila* genome, intergenic lncRNAs were paired with PCGs located at a median distance of 2.1 kb, with 36% of pairs located less than 1 kb apart. We categorized the expression profile of each lncRNA according to its changes over time from L3 to EP and LP, classifying them into 4 groups: increasing, decreasing, peak or valley (Fig. 2B; Supplemental Fig. S1F,G). The same categorization was applied to their associated PCGs. Each lncRNA-PCG association was then classified as concordant if the classification of both genes matched, discordant if their classification was opposite (increasing-decreasing and peak-valley), and unrelated if they were not concordant or discordant or if the associated PCG was not expressed at any stage. The proportions of concordant and discordant pairs were quite similar across the tissues (Fig. 2C-C’’) (Supplemental Table S1), and we found a significant positive correlation for lncRNA-PCG concordant pairs and a significant negative correlation for discordant pairs (Fig. 2D-D’’).

**Figure 2.**
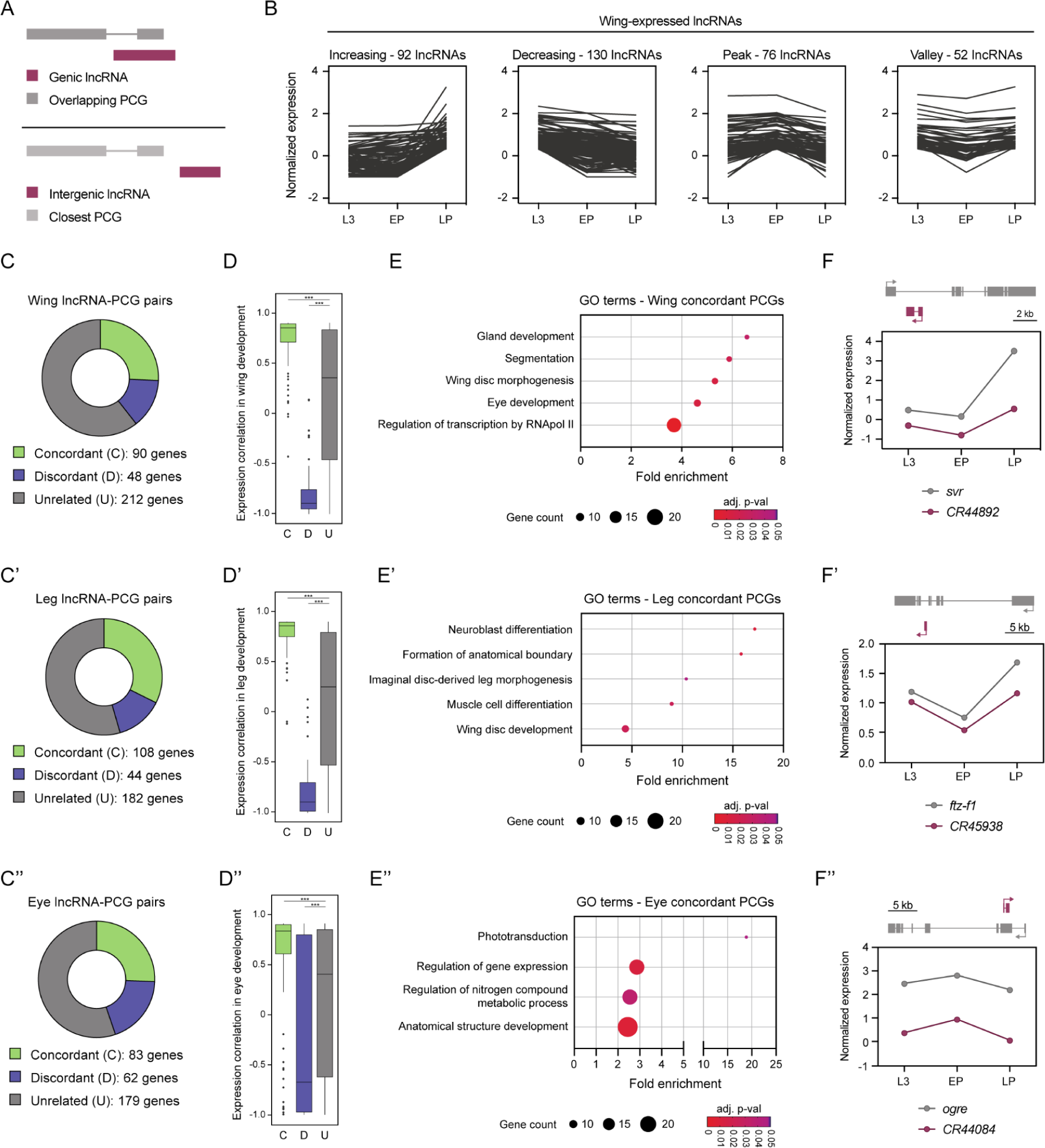
Association of lncRNAs with PCGs in the developing wing, leg and eye discs. (A) Association of genic and intergenic lncRNAs with their overlapping and closest PCGs, respectively. (B) Classification of the expressed lncRNAs in the wing disc according to their developmental expression profile. (C) Classification of all the lncRNA-PCG pairs in the wing (C), leg (C’) and eye (C’’) discs. (D) Pairwise Pearson correlation coefficient of concordant, discordant and unrelated lncRNA-PCG pairs in the wing (D), leg (D’) and eye (D’’) discs. (E) Gene Ontology (GO) term enrichment analysis of the PCGs from the concordant lncRNA-PCG pairs in the wing (E), leg (E’) and eye (E’’) discs. (F) Examples of concordant lncRNA-PCG pairs from the wing disc morphogenesis term (F), imaginal disc-derived leg morphogenesis term (F’), and phototransduction term (F’’). The genomic positioning of the lncRNA and its associated PCG is represented. (***) *p* < 0.001.

We performed a Gene Ontology (GO) term enrichment analysis of the concordant PCGs. In the wing disc, we observed an enrichment in genes involved in development, including segmentation and wing disc morphogenesis (Fig. 2E). For instance, the coding gene *silver* (*svr*), which contributes to the proper formation of the adult wing in terms of shape and size (Sidyelyeva et al. 2006), showed a concordant developmental profile with the lncRNA *CR44892*, which is in an antisense orientation within the first intron of *svr* (Fig. 2F). Concordant PCGs from the leg disc were enriched in several development-related processes, such as neuroblast and muscle cell differentiation, as well as imaginal disc-derived leg morphogenesis (Fig. 2E’). The PCG *ftz transcription factor 1* (*ftz-f1*), which participates in leg development in response to steroid hormone activation (Rewitz et al. 2010), showed a comparable expression profile to the intronic lncRNA *CR45938* (Fig. 2F’). In the eye disc, we found an enrichment in phototransduction-related genes, which are required for the proper development of the eye disc (Fig. 2E’’). One of these genes, the PCG *optic ganglion reduced* (*ogre*), participates in the formation of the gap junctions needed for the transduction of light signals from the photoreceptor to the central nervous system (Curtin et al. 2002; Holcroft et al. 2013), and shows a concordant expression profile with the exonic antisense lncRNA *CR44084* (Fig. 2F’’). These concordant lncRNA-PCG associations may reflect either the sharing of gene-regulatory elements, or the action in *cis* of the lncRNAs in the modulation of the expression of the overlapping PCGs.

### Expression of lncRNAs in regenerating wing discs

Since lncRNAs seem to play important roles in cellular responses to stress, we analyzed the expression profiles of lncRNAs in wing imaginal discs after the induction of cell death using the proapoptotic gene *reaper* (*rpr*). We used previously obtained RNA-seq data from different stages of damage recovery: early, mid and late stages corresponding to 0 hours, 15 hours and 25 hours after stopping cell death induction, respectively (Vizcaya-Molina et al. 2018) (Fig. 3A). To obtain a robust list of differentially-expressed (DE) lncRNAs, we performed pairwise comparisons between control and regeneration samples for each time-point. A high-confidence list of 131 DE lncRNAs in regeneration was obtained (Supplemental Table S2). We found 29 upregulated and 33 downregulated lncRNAs in the early stage of regeneration, followed by 11 upregulated and 56 downregulated lncRNAs in the mid stage, and 22 upregulated and 30 downregulated in the late stage (Fig. 3B). The majority of these lncRNAs were DE at one time-point (71.8%, 94 genes), while 9.9% (13 genes) were DE throughout the entire regeneration process (Fig. 3B). Around half of DE lncRNAs were intergenic (48.8%), while genic lncRNAs were located predominantly in exonic regions (29.8%) compared to introns (21.4%) (Fig. 3C). No significant differences were observed between DE and non-DE lncRNAs in terms of intergenic/genic ratio, transcript length or number of exons (Supplemental Fig. S2A-D).

**Figure 3.**
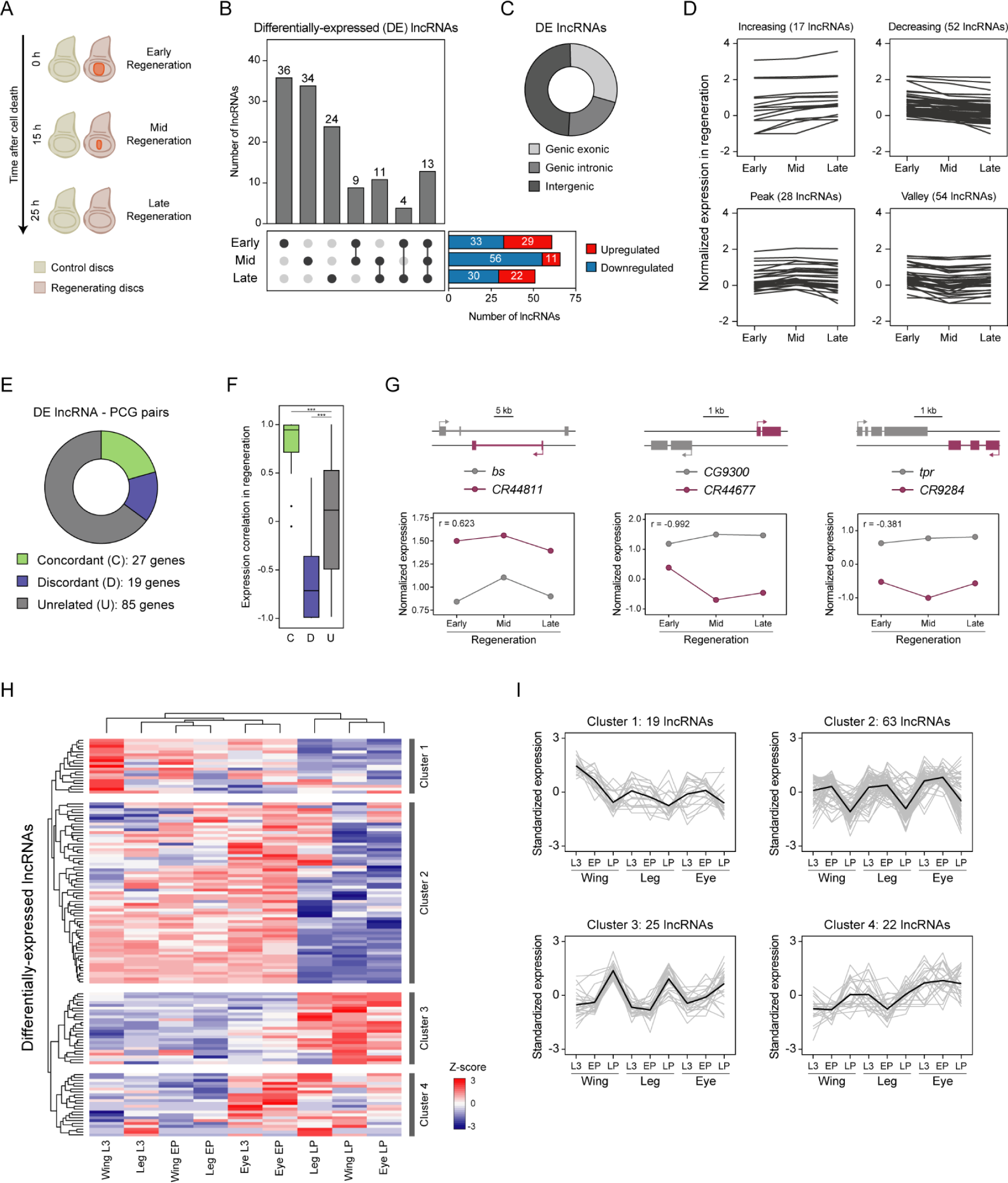
Set of differentially expressed (DE) lncRNAs in regeneration. (A) Representation of the regeneration stages studied in the wing imaginal disc. After cell death induction by expressing *rpr* in the *spalt* domain, 3 different regeneration stages were characterized: early regeneration (0 h after cell death induction; ACD), mid regeneration (15 h ACD) and late regeneration (25 h ACD). (B) Number of upregulated and downregulated lncRNAs at each regeneration stage. Total number of lncRNAs differentially expressed at each time-point or in any combination of time-points. (C) Distribution of DE lncRNAs into genic exonic, genic intronic or intergenic. (D) Classification of the 131 DE lncRNAs in regeneration according to their expression profile during regeneration samples. (E) Percentage of concordant, discordant and unrelated DE lncRNA-PCG pairs. (F) Pairwise Pearson correlation coefficient of concordant, discordant and unrelated DE lncRNA-PCG pairs. (G) Regeneration expression profiles of a concordant lncRNA-PCG pair (left), discordant pair (middle) and unrelated pair (right). (H) Gene expression in the wing, leg and eye disc development of the 131 DE lncRNAs in regeneration. Gene expression is normalized to Z-score values. Complete hierarchical clustering was used to identify the gene clusters. (I) Standardized expression profile of each lncRNA from each cluster. Mean expression profile is represented in black. (***) *p* < 0.001.

Next, we classified the 131 DE lncRNAs according to their expression profile across time in regeneration (Fig. 3D), and associated each of them with their overlapping or closest PCG. This resulted in 131 lncRNA-PCG pairs (131 unique lncRNAs and 121 unique PCGs), 20.6% of which were concordant, 14.5% were discordant, and 64.9% were unrelated (Fig. 3E; Supplemental Table S3). As expected, we observed a high positive correlation for lncRNA-PCG concordant pairs and a strong negative correlation for discordant pairs compared to unrelated pairs (Fig. 3F). We highlighted the concordant pair composed by the natural antisense transcript *CR44811*, which is known to regulate the isoform usage of the overlapping *blistered* (*bs*) gene (Pérez-Lluch et al. 2020), and the discordant intergenic lncRNA *CR44677* and *CG9300,* which showed a high negative correlation in comparison with the unrelated pair composed by *CR9284* and *tpr* (Fig. 3G).

To better characterize the lncRNAs DE in regeneration, we analyzed their expression during normal development in imaginal discs (Ruiz-Romero et al. 2022). Our analysis identified 4 distinct clusters of lncRNAs: (1) those exhibiting peak expression in the wing disc at the L3 stage, (2) those with decreasing expression at the LP stage, (3) those with increasing expression at the LP stage, and (4) those with increased expression in the eye (Fig. 3H,I). The dynamic expression profiles of DE lncRNAs throughout development indicates a tight regulation and a potential role in development. However, upon extending our analysis to the context of regeneration, we observed that these developmental clusters were not maintained (Supplemental Fig. S2E,F), suggesting a context-dependent regulation of their expression.

### LncRNA *CR40469* is not required for normal development but plays a role during wing regeneration

From the set of 131 DE lncRNAs, we searched for putative candidate genes suitable for generating a mutant. To minimize any interference on overlapping genes, we only considered intergenic lncRNAs. We focused on lncRNAs upregulated during early regeneration, particularly the lncRNA *CR40469* (Fig. 4A). *CR40469* is a monoexonic lncRNA spanning 214 bp and localized in the subtelomeric region of the X chromosome, positioned roughly 120 kb away from the chromosome end. It is generally not expressed throughout development, yet, it can be detected in the L3 wing disc (Graveley et al. 2011; Ruiz-Romero et al. 2022). Based on the criteria from Fig. 3E, *CR40469* was classified as unrelated, as the closest PCG, *CG17636*, is not expressed in regeneration.

**Figure 4.**
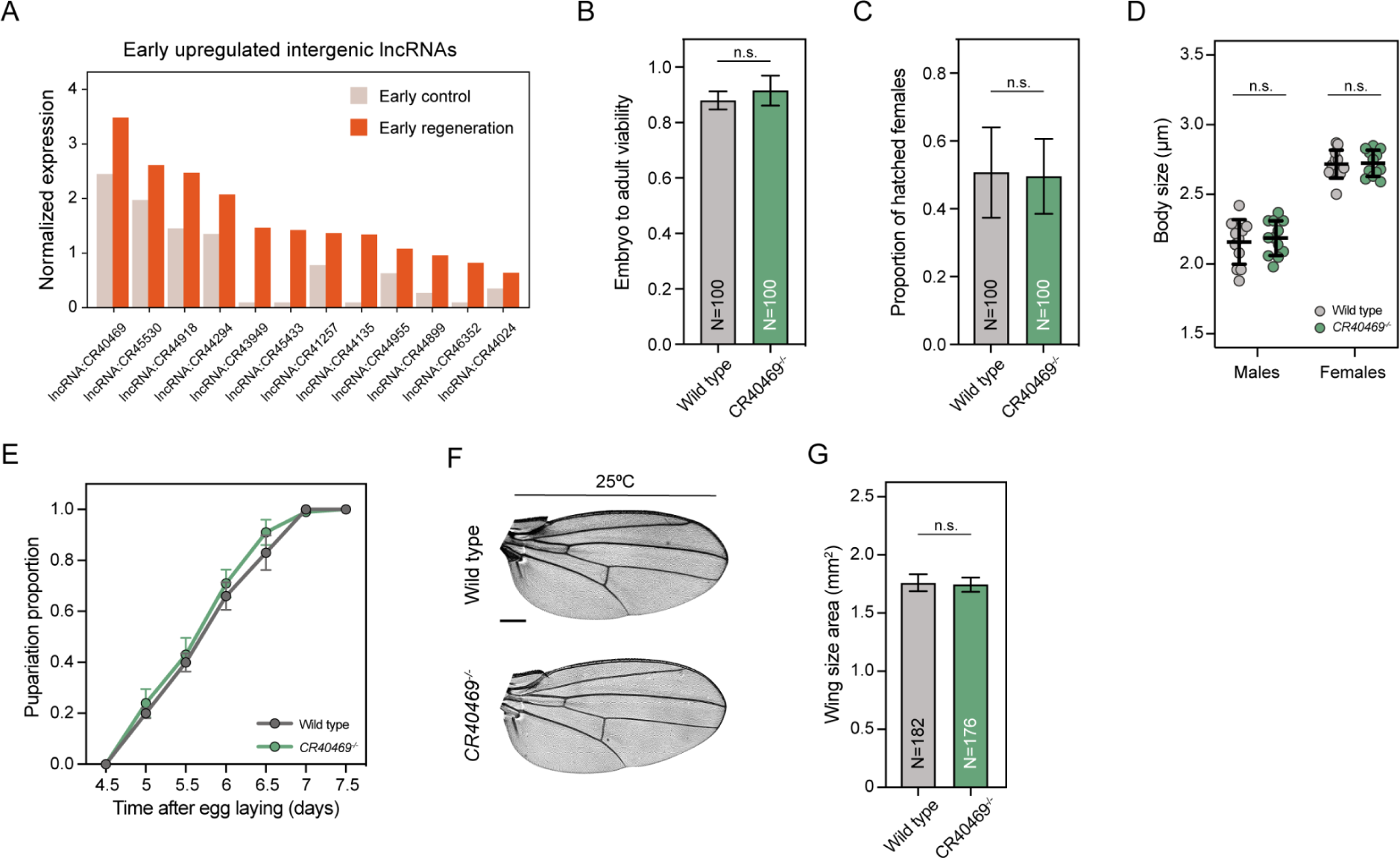
Deletion of *CR40469* does not affect fly development. (A) Expression of the 13 intergenic and early upregulated lncRNAs in early control and early regeneration. (B) Embryo-to-adult viability presented as the proportion of hatched adults from deposited embryos. Four replicates of N=25 each were used. (C) Proportion of hatched females from the emerging adults. Four replicates of N=25 each were used. (D) Adult body size analysis by measuring the length from the head to the abdomen. N=12 per condition. (E) Pupariation analysis showing the proportion of flies that reached the pupal stage at each time-point. Four replicates of N=25 each were used. (F) Sample images of the adult wings of wild type flies and *CR40469* mutants incubated at 25°C. (G) Wing size of the wings represented in F measured as the wing blade area. Scale bar = 250 µm. The mean and standard error of the mean (SEM) are presented in B-C and E. The mean and standard deviation (SD) are presented in D and G. n.s. = non-significant.

We generated a knockout mutant by using the ends-out homologous recombination technique, deleting the entire *CR40469* locus as well as 663 bp upstream and 212 bp downstream (deleting a total region of 1,089 bp), and replacing them with an *mCherry* cassette (Supplemental Fig. S3A-B). Homozygous mutant animals were fully viable (Fig. 4B), reached adulthood at the same sex proportions as the wild type flies (Fig. 4C), and did not show differences in adult body size compared to wild types (Fig. 4D), indicating that the deletion of *CR40469* did not compromise the normal development of male and female flies. We did not find differences in the pupariation time either (Fig. 4E), indicating that mutants did not undergo developmental delays. Also, the deletion of *CR40469* did not affect the adult wing size or morphology (Fig. 4F,G). Thus, we concluded that *CR40469* is not required for fly development under normal laboratory conditions.

Since we identified *CR40469* as a lncRNA upregulated in wing regeneration, we tested the effects of knocking out *CR40469* after tissue damage. For this, we genetically activated the proapoptotic gene *rpr* in the central zone of the wing disc (*spalt* domain). We used the *LHG/LexO* system, which allows conditional inactivation by the temperature sensitive *Gal80^TS^*(Fig. 5A). We did not observe defects in the wing patterning when flies were maintained at permissive temperature (17°C) (Supplemental Fig. S4A). However, after cell death induction, mutant flies showed a significant reduction in their wing regeneration capacity, as there was a higher proportion of aberrant non-regenerated wings characterized by major disruptions in the vein patterning, such as the absence of complete sets of veins and crossveins as well as the appearance of notches in the margins of the wing blade (Fig. 5B,C).

**Figure 5.**
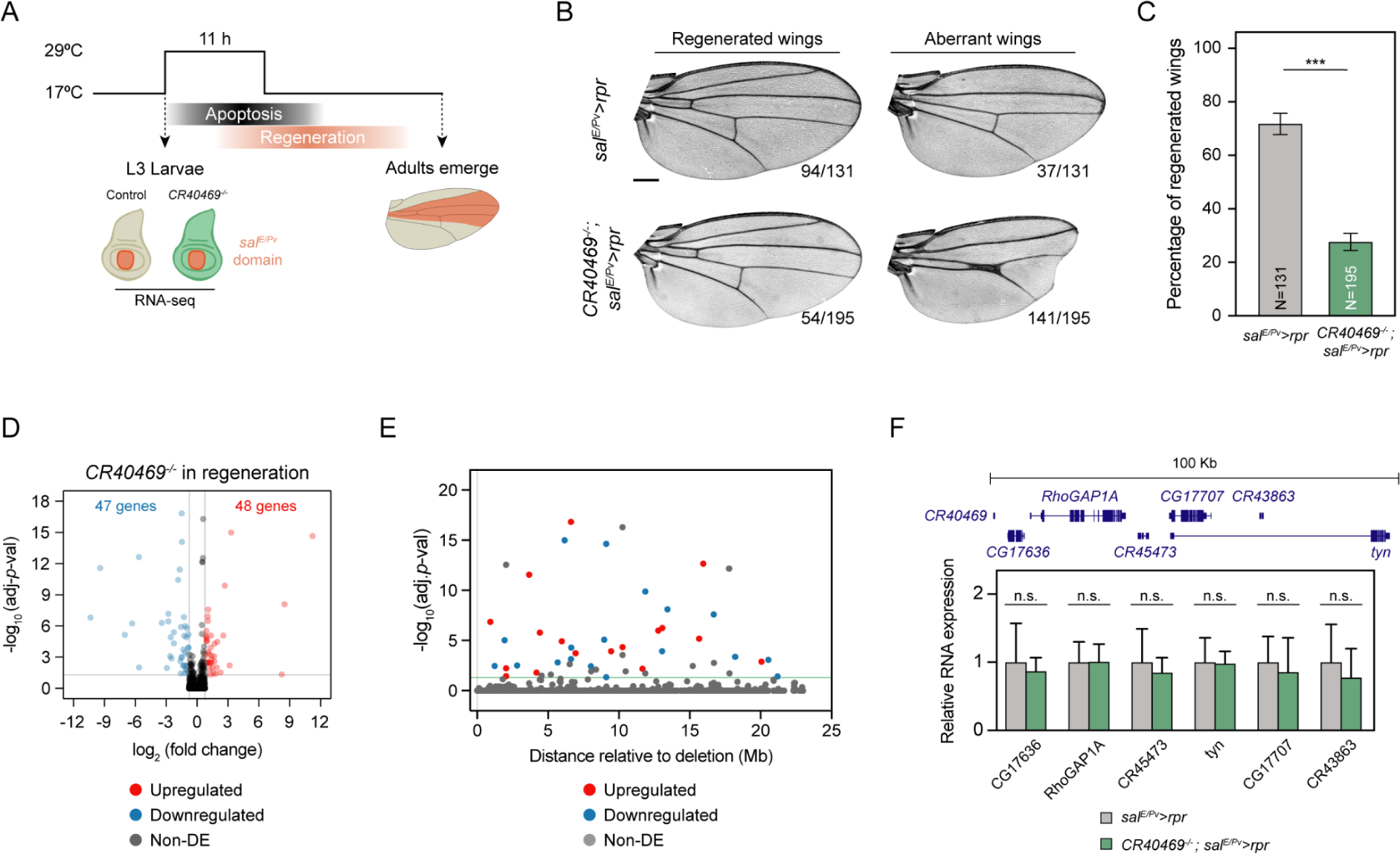
Wing regeneration is impaired in the *CR40469* mutants. (A) Schematic representation of the method used to analyze the regeneration capacity of flies. Cell death was induced in the *spalt* domain of the wing disc for 11 h at the L3 stage, then flies were allowed to recover until adulthood. RNA-seq of control and *CR40469* homozygous mutants were performed following cell death induction. (B) Sample images of most common phenotypes of regenerated and aberrant wings in control and *CR40469* mutants. The number of wings per phenotype is represented. Scale bar = 250 µm. (C) Percentage of regenerated wings from B. (D) Volcano plot displaying the upregulated (red), downregulated (blue) and non-DE genes (gray) in regenerating mutants compared to regenerating controls. (E) Distances of all genes located on the X chromosome relative to the position of *CR40469*. Upregulated, downregulated and non-DE genes are shown in red, blue and gray, respectively. (F) Expression analysis by qPCR of genes located < 100 kb from the *CR40469* locus in the wild type and homozygous mutant wing discs. 3 biological replicates are measured. The mean and SD are represented in C and F.

To characterize the molecular changes occurring in *CR40469* mutants after cell death induction in the wing discs, we analyzed by RNA-seq the gene expression profiles at the early stage of regeneration. Non-mutant regenerating discs were used as controls. We found a total of 95 DE genes after cell death induction in the *CR40469* mutants: 48 of which were upregulated and 47 downregulated (Fig. 5D; Supplemental Table S4). To determine whether the deletion of *CR40469* had a local effect on gene expression, we mapped the genomic position of the DE genes along the X chromosome. The 112 genes located up to 908 kb downstream of the deleted region showed no significant changes in gene expression, with the closest upregulated and downregulated genes located 1.23 Mb and 923 kb away from the *CR40469* locus, respectively (Fig. 5E). We next conducted qPCR analysis to examine the expression of genes located within a 100 kb vicinity of the deletion site. Our results confirmed those obtained from the RNA-seq analysis. Specifically, none of the genes analyzed showed altered expression in regenerating *CR40469* mutant wings compared to the wild type flies (Fig. 5F). Thus, we concluded that the absence of *CR40469* had no significant impact on the expression of nearest genes during regeneration.

### CR40469 is duplicated in the Drosophila melanogaster genome

To further characterize *CR40469*, we performed a BLAST search to assess its putative conservation, but we could not identify a similar sequence in any other species, including other *Drosophila* species. However, we identified the entire *CR40469* genomic sequence with 99.1% similarity (212/214 identities) within the annotated lncRNA *CR34335* sequence in the *Drosophila melanogaster* genome (Fig. 6A; Supplemental Fig. S3C). *CR34335* is located 3.2 Mb downstream of the *CR40469* locus in the X chromosome and is positioned inside a long intron of the PCG *Dpr-interacting protein a* (*DIP-a)*, which is not expressed in development or regeneration in the wing disc (Ruiz-Romero et al. 2022). The *CR34335* gene is monoexonic and contains two polyadenylation signals (Fig. 6A), giving rise to two predicted transcripts that span 249 and 361 nucleotides. Both isoforms contain the entire sequence identical to *CR40469*. Despite their sequence similarity, *CR40469* and *CR34335* exhibit opposite expression patterns. In the development of wing, leg, and eye discs, *CR40469* shows minimal expression, while *CR34335* is highly expressed (Fig. 6B). Conversely, during wing disc regeneration, *CR40469* is upregulated at early and late stages, whereas *CR34335* is consistently downregulated across all time points (Fig. 6C). Using the Multiz alignment tool (Blanchette et al. 2004), we explored the conservation of both lncRNAs in other *Drosophila* species. No traces of synteny nor sequence homology were found for the *CR40469* locus, however, small regions of high similarity were detected for *CR34335* (Fig. 6D). Particularly, we highlighted two conserved blocks located at the edges of the *CR34335* sequence shared with *CR40469* (Fig. 6D). The high conservation of these regions in evolutionarily close *Drosophila* species, the absence of conservation of the *CR40469* locus, and the high sequence identity of *CR34335* and *CR40469* in *Drosophila melanogaster*, suggest that the *CR40469* lncRNA might have originated from the *CR34335* locus specifically in the *Drosophila melanogaster* species.

**Figure 6.**
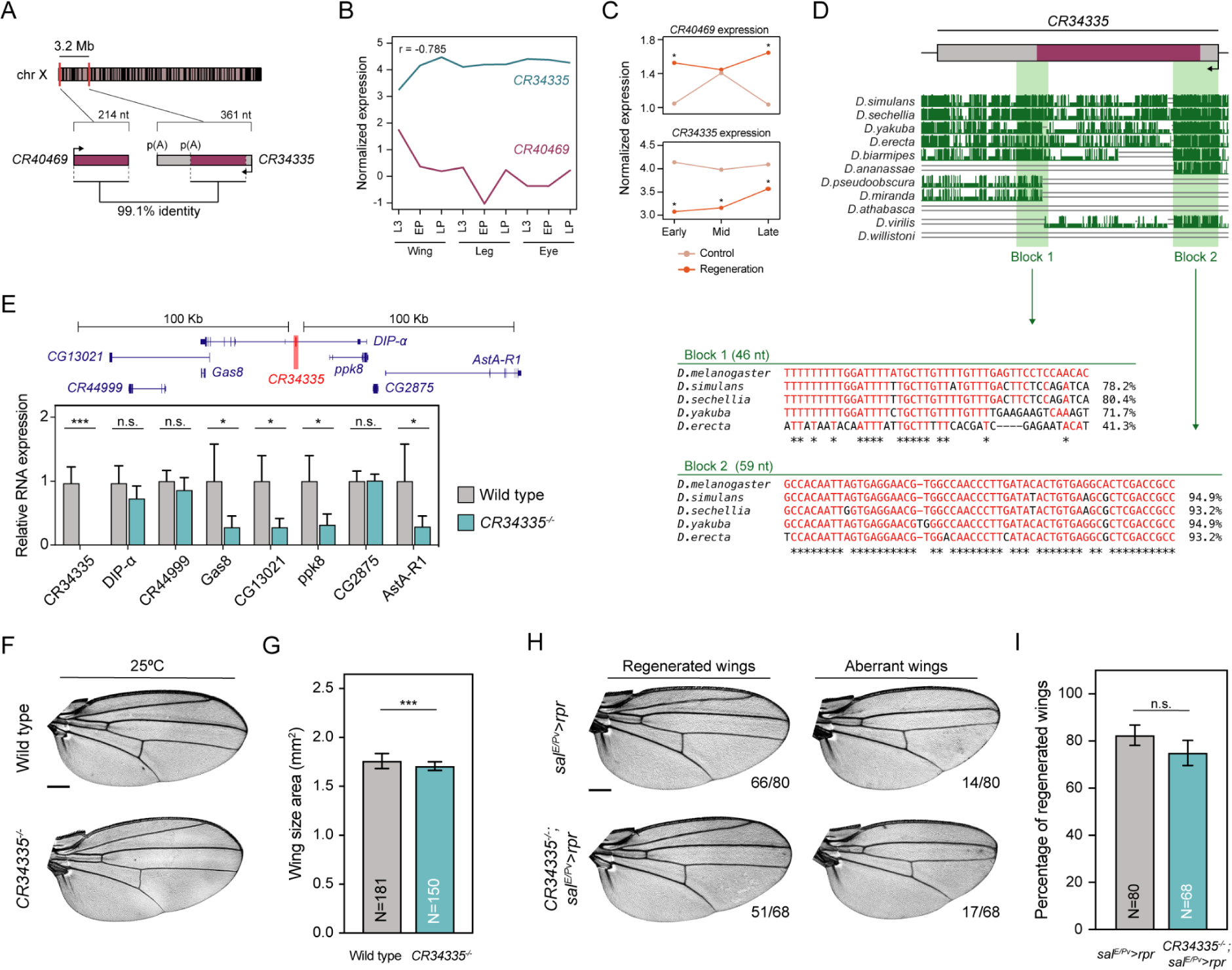
The sequence of *CR40469* is duplicated in the *Drosophila* genome. (A) Schematic representation showing the genomic position of the lncRNAs *CR40469* and *CR34335*. (B-C) Expression profile of *CR40469* and *CR34335* in development and regeneration. Normalized expression is represented as log_10_(TPMs+0.1). Pairwise Pearson correlation coefficient of both lncRNAs is shown. (D) Multiz alignments of the *CR34335* locus in different *Drosophila* species. The genomic sequence of blocks 1 and 2 are represented, as well as their sequence identity compared with the *Drosophila melanogaster* sequence. (E) Expression of *CR34335* and the genes located < 100 kb away in the control and *CR34335* mutant wing discs measured by qPCR. 3 biological replicates per condition are measured. (F) Sample images of the adult wings of wild type flies and *CR34335* mutants incubated at 25°C. (G) Wing size of the wings represented in F measured as the wing blade area. (H) Sample images of most common phenotypes of regenerated and aberrant wings in control and *CR34335* mutants. The number of wings per phenotype is represented. (I) Percentage of regenerated wings from H. The mean and SD are represented in E, G and I. Scale bars in F and H = 250 µm. (***) *p* < 0.001; (*) *p* < 0.05; n.s. = non-significant.

We next analyzed an available *CR34335* mutant that had an insertion of a 7-kb transposon within its exonic sequence, affecting both isoforms. To understand the impact of the transposon insertion, we examined by qPCR the expression of *CR34335* and the 7 genes located up to 100 kb upstream and downstream of the insertion site. We confirmed the absence of *CR34335* expression in the *CR34335* mutant and noted the downregulation of 4 nearby genes (Fig. 6E). Homozygous mutant flies were fully viable and, despite a slight reduction in size (∼3%) compared to wild type wings (Fig. 6F,G), we did not detect any abnormality in the shape or vein patterning (Fig. 6F).

To test whether *CR34335* is necessary for wing regeneration, we used the same system as described above to transiently activate *rpr* in the *spalt* domain of the wing disc. At permissive temperature, mutants developed normal wings (Supplemental Fig. S4A). Upon *rpr* activation, we did not observe a significant decrease in the regeneration capacity of *CR34335* mutants compared to controls (Fig. 6H,I), indicating that *CR34335* is dispensable for regeneration.

Finally, since subcellular localization is one of the primary factors determining lncRNA function, we performed fluorescence *in situ* hybridization (FISH) using a probe complementary to both lncRNAs (*214-probe*). A strong signal was found in the cytoplasm of wild type discs, and it was even stronger in the *CR40469* mutants, indicating that *CR34335* is a cytoplasmic lncRNA (Supplemental Fig. S5A-D). Nevertheless, probably due to its lower expression levels in development, we could not detect *CR40469* transcripts in the wing imaginal disc (Supplemental Fig. S5E,F).

### Regeneration mutant phenotypes are rescued by ectopic expression of *CR40469*

To elucidate whether the ectopic expression of *CR40469* or *CR34335* was sufficient to restore the regeneration capacity of *CR40469* homozygous mutants, we generated transgenic flies carrying a copy of the *CR40469* or the *CR34335* genes downstream of a UAS sequence. To assess the efficiency of the transgenes, we performed *in situ* hybridization using a *nubbin* driver (*nub-Gal4*) to induce their expression in the pouch of the wing disc in a *CR34335* homozygous mutant background. This choice was made to avoid the potential masking effect of high *CR34335* levels on the detection of the ectopic expression of the transgenes. Similar to the endogenous *CR34335* (Supplemental Fig. S5A-D), we observed ectopic *CR34335* also in the cytoplasm (Fig. 7A). Also, upon inducing *CR40469* expression in *CR34335* mutants, a cytoplasmic signal was detected (Fig. 7A), suggesting that the endogenous *CR40469* might also be located in the cytoplasm. We next analyzed the effects of the ectopic expression of *CR40469* and *CR34335* in the wing in a wild type background using *nub-Gal4*, whose domain encompasses the entire wing blade in the adult. In both cases, adult wings showed normal vein patterning (Fig. 7B) and size (Fig. 7C), indicating that the overexpression of *CR40469* or *CR34335* does not affect normal wing development. Similarly, no visible defects were detected in adult flies when using an ubiquitous driver (data not shown). Next, we tested whether expression of *CR40469* or *CR34335* could rescue the aberrant wing phenotypes observed upon the activation of cell death in *CR40469* homozygous mutants. We combined the induction of cell death in the *spalt* domain with the expression of *CR40469* or *CR34335* in the *nub* domain. The ectopic expression of *CR40469* significantly increased the proportion of regenerated wings observed in *CR40469* mutant flies following the induction of cell death (Fig. 7D,E). Additionally, *CR40469* expression led to significantly larger wings compared to controls, indicating that not only vein patterning but also wing size is recovered (Fig. 7F). On the other hand, expressing *CR34335* in the *nub* domain of *CR40469* mutants also resulted in a higher number of regenerated wings, although wing size was not significantly restored (Fig. 7D-F). Taken together, our findings point to a potential *trans*-acting role of *CR40469* during the damage response, with *CR34335* potentially performing a partially redundant function in regeneration when *CR40469* is absent.

**Figure 7.**
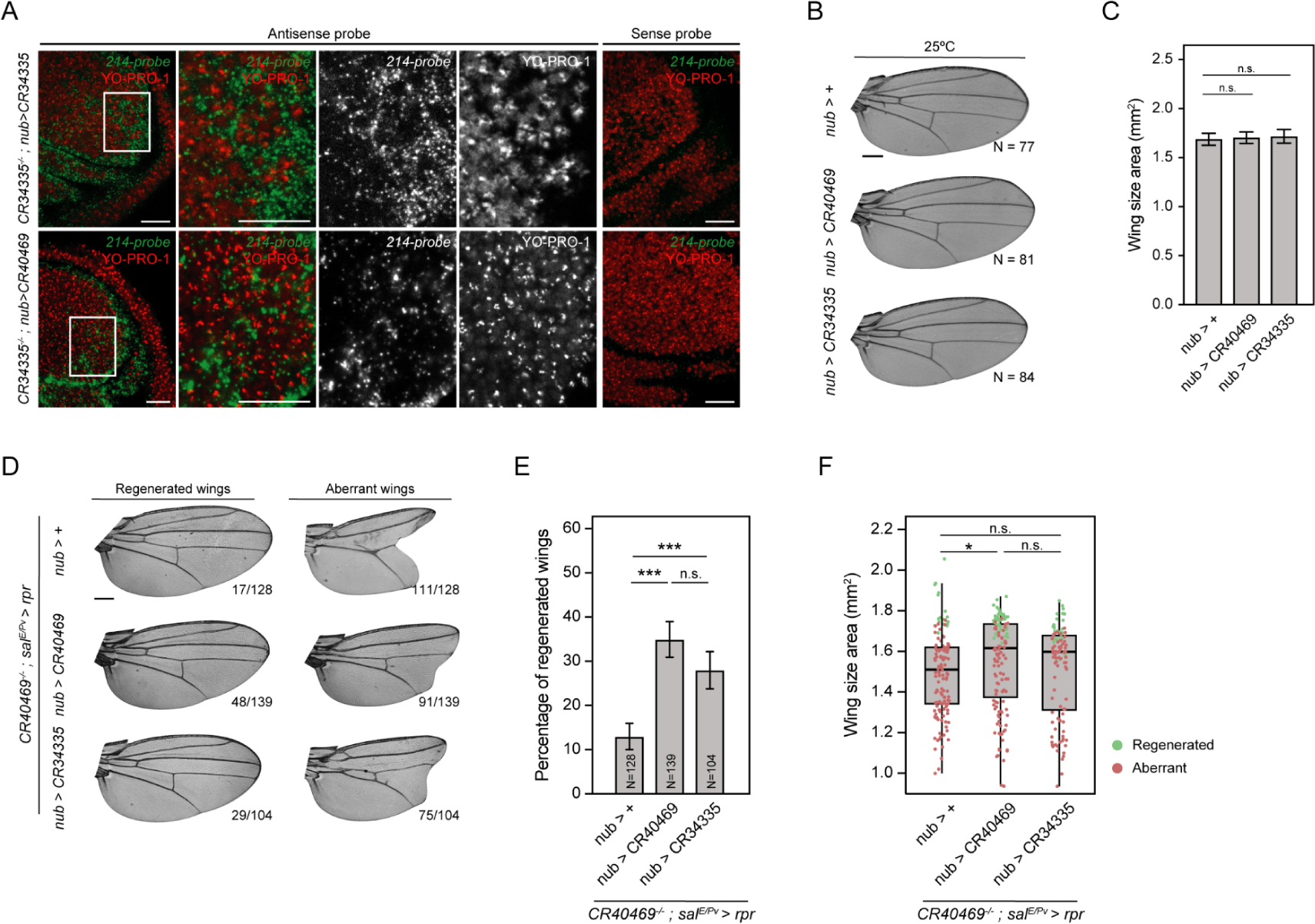
Ectopic expression of *CR40469* rescues the *CR40469* mutant regeneration phenotypes. (A) *In situ* hybridization of the wing imaginal disc after expression of *CR34335* (top row) or *CR40469* (bottom row) driven by a *nub* promoter in a *CR34335* homozygous mutant background. *214-probe* is complementary to both *CR40469* and *CR34335*. N ≥ 4 per condition. Scale bar = 20 µm. (B) Sample wings of *nub>+* (control), *nub>CR40469* and *nub>CR34335* incubated at 25°C. (C) Wing size of genotypes represented in B. (D) Sample images of most common phenotypes of regenerated and aberrant wings of *nub>+*, *nub>CR40469* and *nub>CR34335* in a *CR40469* homozygous mutant background. The number of wings per phenotype is represented. (E) Percentage of regenerated wings from D. (F) Wing size of genotypes represented in D. Quartiles, interquartile range and median are represented. Regenerated and non-regenerated wings are shown in green and red, respectively. The mean and SD are represented in C and E. Scale bars in B and D = 250 µm. (***) *p* < 0.001; (*) *p* < 0.05; n.s. = non-significant.

## Discussion

Non-coding transcripts are being increasingly identified in transcriptional studies and there is growing evidence of lncRNA functionality. It is likely that the most expressed lncRNAs are actively or potentially involved in several cellular processes (Lee et al. 2019). Here, with the aim of describing the participation of lncRNAs in development and regeneration, we have studied their expression in *Drosophila* imaginal discs. Compared to the around 8,000 PCGs expressed in the discs, we have only detected the expression of around 200 lncRNAs. As previously reported for *Drosophila* and other species (Derrien et al. 2012; Brown et al. 2014; Washietl et al. 2014), the tissue and stage specificity of lncRNAs is remarkably higher than that of PCGs regardless of the tissue or developmental stage analyzed, implying a tight control of lncRNA transcription in the developing imaginal discs.

One of the most studied functions of lncRNAs is their ability to influence gene expression at multiple levels, including the modulation of chromatin accessibility, the activation of RNA polymerase II, the recruitment of transcription factors, and the regulation of mRNA elongation (Statello et al. 2021). LncRNAs can act in *trans*, functioning far and independently of their transcription site (Lewandowski et al. 2019; Ariel et al. 2020), or in *cis*, acting near and depending on their locus (Gil and Ulitsky 2020). In humans, the expression of lncRNAs tends to correlate with the expression of overlapping and genomically close PCGs (Derrien et al. 2012). Currently, there is no similar data in *Drosophila*, but the higher genome compactness may reinforce the same idea. Here, we have associated each lncRNA expressed in the developing discs with their overlapping or closest PCG. Regardless of the tissue type, ∼25% of the analyzed pairs have shown concordant expression profiles in development. We postulate that some of these concordant lncRNAs could regulate the expression of their associated PCG in *cis*. Consequently, we have analyzed the functions of the concordant PCGs to infer the biological processes in which their paired lncRNAs might participate. Our findings highlight that lncRNAs expressed in the developing discs are preferentially located near PCGs involved in key developmental processes, such as wing disc morphogenesis in the wing disc, phototransduction in the eye disc, and imaginal disc-derived leg morphogenesis in the leg disc. Although we cannot discard the fact that some of the concordant lncRNA-PCG pairs might just reflect the sharing of the same regulatory elements, some of these lncRNAs could be involved in the regulation of key PCGs during development.

Evaluating the functionality of lncRNAs presents significant challenges, and the effects of their lack of function can only be determined conclusively after thorough monitoring of the potential phenotypic impact across various conditions. We chose to study lncRNAs expression and function following damage because of the dynamic changes that take place during the process of regeneration. A few studies have undertaken global analyses of lncRNAs in regeneration, for instance following injury in skeletal muscle (Gonçalves et al. 2017) or during cardiac regeneration in zebrafish (Lumley et al. 2021). Specific lncRNAs have also been linked to regeneration: H19 is associated with nerve degeneration and regeneration in rats (Li et al. 2022), and mucosal regeneration in mice (Geng et al. 2018); lncMREF is a positive regulator of muscle regeneration in mice, pigs and humans (Lv et al. 2022); *CR46040* has recently been identified as crucial for the proliferation of intestinal stem cells in response to injury in *Drosophila* (Xu et al. 2023). In the context of wing regeneration, we have identified a subset of 131 DE lncRNAs that could participate in the recovery process. In line with their high specificity, most lncRNAs are DE in one particular time-point, suggesting that they might be required only during specific periods. In contrast to the peak of active transcription described for PCGs (Vizcaya-Molina et al. 2018), DE lncRNAs tend to be downregulated in regeneration rather than upregulated, suggesting that coding and non-coding genes are regulated differently in response to damage.

We have focused our study on the intergenic lncRNA *CR40469,* which is upregulated after damage in the wing imaginal disc and is required for the regeneration process. *CR40469* is exclusively found in *Drosophila melanogaster* and its sequence is included within the sequence of the lncRNA *CR34335*, whose 5’ and 3’ ends are conserved in a variety of *Drosophila* species. This led us to hypothesize that *CR40469* may have emerged from the *CR34335* locus through a duplication event specific to *Drosophila melanogaster*. In fact, *CR40469* is located in the subtelomeric region of the X chromosome, immediately after the most proximal *HeT-A* element, one of the retrotransposons responsible for telomere elongation in flies (George et al. 2006; Casacuberta 2017), reinforcing the hypothetical scenario of retrotransposition involving *CR34335*, which is located 3.2 Mb downstream. Both genes show an opposite expression pattern: while *CR40469* is probably silenced within heterochromatin in most cells and conditions, *CR34335* is highly and ubiquitously expressed at similar levels of genes encoding for ribosomal subunits (Supplemental Fig. S6A,B). Upon cell death, however, *CR40469* is upregulated and *CR34335* is downregulated. This contrasting expression profile, coupled with their high sequence similarity, suggests that these lncRNAs might engage in competitive binding for specific factors involved in regulating their expression or stability.

The subcellular localization of lncRNAs is crucial in determining their molecular function (Carlevaro-Fita et al. 2019). We have detected *CR34335* transcripts in the cytoplasm. Although we could not detect endogenous *CR40469* transcripts, we have identified ectopic *CR40469* also in the cytoplasm, indicating that both lncRNAs are exported from the nucleus. Multiple studies have indicated that the majority of cytoplasmic lncRNAs localize to ribosomes (van Heesch et al. 2014; Carlevaro-Fita et al. 2016). It is speculated that some of these lncRNAs may be translated into small peptides (Ruiz-Orera et al. 2014), some of which have been shown to be functional (Galindo et al. 2007; Magny et al. 2013; Nelson et al. 2016). Indeed, a short peptide consisting of 33 amino acids is predicted from both *CR40469* and *CR34335* (Supplemental Fig. S6C). While *CR34335* transcripts have been previously associated with ribosomes in early embryos, the putative translation of the small peptide has not been detected (Li et al. 2016). Other functions attributed to ribosome-bound lncRNAs include the regulation of mRNA translation and stability (Carlevaro-Fita et al. 2019), which could be relevant to the roles of *CR40469* and *CR34335*. Moreover, it is important to consider the influence of the extra nucleotides at the 5’ and 3’ ends of *CR34335* compared to *CR40469*, which might result in diverse secondary structures or binding associations, thus potentially affecting their molecular function.

In addition to its subcellular localization, a *trans*-acting role for *CR40469* in regeneration may be implied by the seemingly stochastic distribution of differentially-expressed genes in *CR40469* mutants across the *Drosophila* genome. The restoration of the regeneration capacity in *CR40469* mutants following ectopic expression of *CR40469* further supports its *trans*-acting role. Moreover, the observation that ectopic expression of *CR34335* is able to partially rescue the regeneration capacity of *CR40469* mutants suggests a potential functional substitution of *CR34335* in the absence of *CR40469*. In conclusion, our study identifies *CR40469* as a *trans*-acting cytoplasmic lncRNA participating in wing regeneration. Additionally, as the non-coding genome remains largely unexplored, the partial duplication of lncRNAs that we have uncovered in our study may not necessarily be an exceptional phenomenon, but an instance of a more general mechanism by means of which lncRNAs acquire novel functions.

## Methods

### Drosophila strains

The *Drosophila melanogaster* strains *sal^E/Pv^-LGH* and *lexO-rpr* were previously described (Santabárbara-Ruiz et al. 2015). The following strains were provided by the Bloomington *Drosophila* Stock Center: *Canton S* (RRID: BDSC_64349), *tub-Gal80^TS^* (RRID: BDSC_7017), *nub-Gal4* (RRID: BDSC_86108) and *CR34335^-/-^* (RRID: BDSC_23825). *Canton S* was used as the wild type strain.

### Expression and specificity during development

RNA-seq samples from the wing, eye and leg discs of third instar larvae (L3, 110-115 h after egg laying), early pupae (120-130 h), and late pupae (225-235 h) obtained from Ruiz-Romero and colleagues (Ruiz-Romero et al. 2022) (ArrayExpress accession number: E-MTAB-10879) were used to identify the expressed genes. Genes and transcripts were quantified in transcripts per kilobase million (TPMs) using RSEM v1.2.21 (Li and Dewey 2011). The average TPMs for each gene in each sample was calculated as the average TPMs for two biological replicates. Only genes with an expression of at least 1 TPM were considered expressed. Genes aligned to non-canonical chromosomes were discarded. A total of 13,957 protein-coding genes and 2,455 lncRNAs were considered (FlyBase genome annotation version r6.29 (Attrill et al. 2016; Larkin et al. 2021)).

Genes were considered tissue-specific if their expression was at least 1 TPM in only one particular tissue, independent of the developmental stages in which they were expressed. Stage-specific genes had an expression of at least 1 TPM in only one particular developmental stage, independent of the tissues in which they were expressed.

### LncRNA classification and association with protein-coding genes (PCGs)

LncRNAs were classified with respect to their genome location using the classification module of the FEELnc pipeline (Wucher et al. 2017). FEELnc received the 2,455 annotated lncRNAs from the FlyBase genome annotation version r6.29 as input, classifying the lncRNAs into three broad groups: lncRNAs not overlapping with any other PCGs were considered intergenic, lncRNAs located within an intron of a PCG were classified as genic intronic, and lncRNAs overlapping by at least 1 bp with an exonic sequence of a PCG were classified as genic exonic. The classification was mutually exclusive in the following rank: genic exonic > genic intronic > intergenic.

Intergenic lncRNAs were associated with the closest PCG by measuring the end-to-end distance, independent of the expression, orientation, and direction of transcription. Genic intronic and genic exonic lncRNAs were paired with their overlapping PCGs. For lncRNAs overlapping with multiple PCGs, we only considered the gene showing more overlapping nucleotides.

### Expression profile and lncRNA-PCG pair classification

The expressed lncRNAs shown in Fig. 1B and their associated PCGs (Supplemental Table S1) were each classified as increasing (lowest expression in L3 and highest expression in LP), decreasing (lowest expression in LP and highest expression in L3), peak (highest expression in EP) or valley (lowest expression in EP). The differentially expressed lncRNAs in regeneration shown in Fig. 3B and their associated PCGs (Supplemental Table S3) were each classified according to their expression during regeneration as increasing (lowest expression at the early stage and highest expression at the late stage), decreasing (lowest expression at the late stage and highest expression at the early stage), peak (highest expression at the mid stage) or valley (lowest expression at the mid stage).

LncRNA-PCG pairs were then classified as concordant (if the classification of both members in the pair matched: increasing/increasing, decreasing/decreasing, peak/peak and valley/valley), as discordant (if the classification of both members in the pair was opposite: increasing/decreasing and peak/valley), or as unrelated (if the PCG was not expressed or if they were not classified as concordant or discordant: increasing/peak, increasing/valley, decreasing/peak and decreasing/valley).

We used R to calculate the pairwise Pearson correlation coefficient (PCC) for each lncRNA-PCG pair. Normalized expression data (log_10_-transformed TPMs plus a pseudocount of 0.1) of the wing disc (Fig. 2D), leg disc (Fig. 2D’), eye disc (Fig. 2D’’) or regeneration samples (Fig. 3F) was used for the PCC calculation.

### Gene Ontology (GO) analysis

The GO enrichment analysis tool (Mi et al. 2019) from the PANTHER 17.0 database was used to determine the GO term enrichment, using biological process trees. Input gene sets were the PCGs from concordant lncRNA-PCG pairs. *p*-values were adjusted using the false discovery rate correction for multiple comparisons.

### Differential expression analysis in regenerating discs

Differential gene expression analyses of the control and regeneration samples, obtained from Vizcaya-Molina and colleagues (Vizcaya-Molina et al. 2018) (GEO Accession number: GSE102841), were performed separately at each time-point. We removed all the genes expressed < 1 TPM in all samples. We used a simple fold change approach of ≥ |1.7| and a DESeq2 (Love et al. 2014) analysis considering an absolute fold change of ≥ 1.7 and a Benjamini-Hochberg adjusted *p*-value < 0.05. Genes DE using both methods were considered. Protein-coding genes *rpr* and *Gadd45* were used as positive controls.

We obtained an initial list of 201 DE lncRNAs. The absence of strand information in these RNA-seq samples hampered the correct attribution of reads between the exonic lncRNAs and their overlapping PCGs. For this reason, we manually validated the initial 109 DE exonic lncRNAs using the RNA-seq bigwig tracks of the UCSC Genome Browser. After manual curation, we removed 70 exonic lncRNAs, ending up with a robust list of 131 DE lncRNAs (Supplemental Table S2). Among them, we found 5 stable intronic sequence RNAs (sisRNAs) and 3 hairpin RNAs (hpRNAs), which were not considered for the selection of our candidate lncRNA.

Due to the 99.1% similarity between the *CR40469* and *CR34335* sequences, we visually inspected the reads mapped to the *CR40469* locus to confirm that mapping was correct. We corroborated that the two mismatched nucleotides between both lncRNAs were sufficient to correctly map the reads to each loci.

### Developmental expression profile of PCGs and lncRNAs

The expression profile of PCGs (Fig. 1E), lncRNAs (Fig. 1F) and differentially-expressed lncRNAs (Fig. 3H) in developing wing, leg and eye discs was represented as a heatmap. Gene expression values were standardized into Z-score values and normalized into a −3 to 3 scale. Gene and sample clustering was performed using a complete hierarchical clustering. Major gene clusters were highlighted for each heatmap.

### Generation of *CR40469* knock-out mutants and UAS transgenic flies

The deletion of *CR40469* was performed using the ends-out homologous recombination technique. Long homology arms for the upstream and downstream regions of the *CR40469* locus were designed. A 1,089-bp containing the entire *CR40469* gene, as well as the upstream 663 bp and downstream 212 bp was excised and replaced by an *mCherry* cassette. The absence of the *CR40469* locus was confirmed by PCR. The designed primer sequences are shown in Supplemental Table S5.

For the ectopic activation of *CR40469* and *CR34335*, we used a semi-directed cloning protocol to clone the entire *CR40469* or *CR34335* genomic sequences into a pUAST plasmid. We designed primers to amplificate the *CR40469* and *CR34335* genes (Supplemental Table S5). An EcoRI target site was added into the 5’ end of forward primers to improve the cloning efficiency. PCR using the following program: 4’ at 95°C, 40 cycles of 30’’ at 95°C, 30’’ at 61°C and 10’’ at 72°C, and a final step of 2’ at 72°C, was performed to amplify the inserts. PCR products were purified using the MinElute PCR purification kit (Qiagen). Fragments were digested using EcoRI-HF (New England Biolabs) and purified using the MinElute Cleanup kit (Qiagen). pUAST vector was double digested using EcoRI-HF and HpaI (New England Biolabs), then the digested vector was run in an electrophoresis gel, and the band located at a 7.9 kb mark was cut and purified using the EZNA Cycle pure kit (Omega Biotek) Purified digested vector and inserts were ligated in a 1:30 vector/insert ratio using the T4 DNA ligase (New England Biolabs) following manufacturer’s protocol. Then, they were transformed into DH5a competent cells (Invitrogen) and antibiotic-resistant clones were sequenced. *CR40469*- and *CR34335*-containing pUAST vectors were injected into the VK33 position at the FlyORF *Drosophila* Injection Service.

### Embryo-to-adult viability and pupariation assays

Flies aged 3-5 days were allowed to lay eggs on standard medium in plates containing yeast paste for 3 hours at 25°C. The eggs were incubated for 16 hours at 25°C. Then, the unhatched embryos were transferred to vials containing standard medium in four replicates containing exactly 25 embryos each, resulting in a total of 100 embryos per condition.

For the analysis of pupariation, the vials were incubated at 25°C and the number of pupae was counted every 12 hours starting 108 hours after egg laying. The pupariation percentage at each timepoint was calculated according to the total number of larvae capable of transitioning to the next developmental phase.

To assess embryo-to-adult viability, the vials were incubated at 25°C for 15 days and the number of hatched adults per vial was counted. Embryo-to-adult viability was calculated as the proportion of adults hatched from the deposited embryos. The proportion of hatched females was calculated as the percentage of females among all the hatched adult flies.

### Analysis of adult body size and wing phenotypes

Flies aged 3-5 days were allowed to lay eggs on standard medium in vials for 3 hours at 25°C. The eggs were incubated at 25°C until adulthood. Then, the adult flies were placed at-20°C for 5 min for immobilization and images from the lateral view were taken under a microscope. 12 adult flies per condition were imaged. The lengths of the head, thorax and abdomen were summed up to obtain a score for the adult body size.

For the study of wing phenotypes, female flies of appropriate genotypes were selected and stored for at least 24 hours in a 1:2 glycerol:ethanol solution. Then, wings were dissected in water, washed in ethanol, mounted in 6:5 lactic acid:ethanol and analyzed under a microscope. Wing size was determined as the area inside the perimeter of the wing blade. Wings were considered aberrant or non-regenerated when missing complete veins and/or when notches were present in the wing blade.

### RNA extraction, reverse transcription and quantitative PCR (qPCR)

For RNA extraction, 50 wing discs per sample were dissected in Schneider’s medium (Sigma Aldrich). The Quick-RNA Microprep kit (Zymo Research) was used following the manufacturer’s instructions to isolate RNA, then it was incubated with DNase I (Promega) at 37°C for 30 minutes and treated with the RNA Clean and Concentrator-5 kit (Zymo Research). A total of 1 µg of RNA was used as a template for cDNA synthesis using Moloney Murine Leukemia Virus reverse transcriptase (M-MLV RT) (Invitrogen).

Reactions containing FastStart Universal SYBR Green Master (Rox) (Roche) and the appropriate cDNA and primers were run in a 7500 Real-Time PCR System (Applied Biosystems). Samples were normalized to the levels of *sply* and fold changes were calculated using the ddCt method. Three technical replicates were used for each reaction, and three separate biological replicates were collected for each experiment. The designed primer sequences are shown in Supplemental Table S5.

### Induction of cell death and transgene activation

To study regeneration, we used the wing specific *sal^E/Pv^*enhancer to drive the expression of LHG in the cells of the central part of the wing disc (*sal^E/Pv^-LHG*), where the pro-apoptotic construct *lexO-rpr* was activated. The LHG is a modified version of lexA that is suppressible by thermosensitive Gal80 (Gal80^TS^) (Yagi et al. 2010). We used an additional *tub-Gal80^TS^*construct to inhibit the expression of *rpr* at 17°C.

Flies were allowed to lay eggs on standard medium in vials for 6 hours at 17°C. Embryos were incubated at 17°C for 192 hours (8 days), before being transferred to a water bath at 29°C for 11 hours for *rpr* activation. Subsequently, they were returned to 17°C until adulthood. Controls lacking the *lexO-rpr* transgene were treated in parallel. Additional controls with the identical genotypes were incubated at 17°C until adulthood. Wing dissection and handling were performed as described above.

We used *nub-Gal4* to induce the expression of *UAS-CR40469* or *UAS-CR3433*5 in the pouch of wing imaginal discs. For constitutive activation of the transgenes, embryos were cultured at 25°C. *In situ* hybridization analysis was performed after 96 hours (4 days) of incubation, while the analysis of adult wings was performed after 15 days. To investigate the effect of transgene activation in the context of regeneration, we followed the same protocol described above using *nub-Gal4* and *sal^E/Pv^-LHG* in combination with *tub-Gal80^TS^* to prevent the expression of both constructs at 17°C.

### RNA-seq sample collection and RNA isolation

For RNA-seq sample collection, flies were allowed to lay eggs on standard medium in vials for 6 hours at 17°C. The discs were incubated at 17°C for 192 hours (8 days), then the vials were transferred to a water bath at 29°C for 16 hours to activate *rpr* expression. L3 larvae from the following genotypes were selected: *lexO-rpr* ; *sal^E/Pv^-LHG:tub-Gal80^TS^* (control) and *CR40469^-/-^; lexO-rpr* ; *sal^E/Pv^-LHG:tub-Gal80^TS^*(*CR40469* homozygous mutants).

50 wing discs per sample were dissected in cold Schneider’s medium. The Quick-RNA Microprep kit (Zymo Research) was used following manufacturer’s instructions to isolate the RNA. Then, it was incubated with DNase I (Promega) at 37°C for 30 minutes and treated with the RNA Clean and Concentrator-5 kit (Zymo Research). The purity and concentration of the resulting RNA were assessed using Nanodrop (Thermo Fisher Scientific) and Qubit (Invitrogen).

### RNA-seq library preparation and sequencing

For library preparation, 500 ng of total RNA were used for reverse transcription. Ribosomal RNA (rRNA) was depleted by selecting the poly-A transcripts. All libraries were sequenced on an Illumina HiSeq2500 sequencer, using 50-bp paired-end reads. Library preparation and sequencing were performed at the Genomics Unit of the Center for Genomic Regulation (CRG).

### Mapping and assembling pipeline of *CR40469^-/-^* RNA-seq data

Transcriptomic data were processed using the grape-nf pipeline (https://github.com/guigolab/grape-nf). Reads were aligned to the fly genome (dm6) using STAR v2.4.0j (Dobin et al. 2013), with up to 4 mismatches per paired alignment, using the FlyBase genome annotation version r6.29. Only the alignments mapping to ten or fewer loci are reported. Genes and transcripts were quantified in TPMs using RSEM v1.2.21 (Li and Dewey, 2011). GTF version r6.29 contains a total of 16,412 genes: 13,957 PCGs and 2,455 lncRNAs. In our study, lncRNAs were defined as non-coding genes with > 200 bp and aligned to canonical chromosomes.

Quality control of the alignment sequencing data was performed using QualiMap v.2.2.1 (García-Andrade et al. 2012) and Picard v.2.6.0 (http://broadinstitute.github.io/picard). Using QualiMap, we obtained the number of reads, the number of mapped reads, the duplication rate and the GC percentage. We obtained the dropout and GC dropout using Picard. Assessment of the reliability of the replicates was measured with weighted correlation network analysis (WGCNA). WGCNA was implemented with the R package WGCNA v1.69 (Langfelder and Horvath 2008). A cutoff of less than 2 standard deviations from a normal distribution was implemented to use a replicate. The number of mapped reads and the correlation between replicates are provided in Supplemental Table S6 and Supplemental Fig. S7, respectively.

### Filtering pipeline and differential expression analysis of *CR40469^-/-^* RNA-seq data

We used the statistical methods implemented in DESeq2 v1.26.0 (Love et al. 2014). Only genes with an expression of at least 1 TPM in at least one sample were selected for the differential expression analysis. The two-factor with interaction approach was implemented, considering the following design matrix: genotype, condition and genotype∼condition, where the genotype is control or *CR40469*^-/-^ and the condition is regeneration or non-regeneration. All genes with an absolute fold change > 1.7 and a Benjamini-Hochberg adjusted *p*-value < 0.05 were considered differentially expressed.

### Multiple sequence alignment

The alignment of the *CR40469* and *CR34335* transcripts was performed using Clustal Omega (Madeira et al. 2022). We used the unique *CR40469* transcript and the two annotated transcripts of *CR34335* as input. The alignment was run using the default parameters.

### Conservation of *CR40469* and *CR34335* in other *Drosophila* species

To analyze the conservation of *CR40469* and *CR34335* in other *Drosophila* species, we used the Multiz alignments tool of the UCSC Genome Browser (Blanchette et al. 2004). For each block of conservation selected, we extracted the genomic sequence of each species and used Clustal Omega (Madeira et al. 2022) to obtain the multiple sequence alignment. Positions identical to the *Drosophila melanogaster* sequence were represented in red.

### Riboprobe synthesis

To analyze the localization of the *CR40469* and *CR34335* transcripts, we synthesized a digoxigenin (DIG)-labeled RNA probe. For this, we selected the 214-bp sequence of the *CR40469* transcript, which is 99.1% identical to the *CR34335* transcripts, for PCR amplification. We named this probe *214-probe*. gDNA from *Canton S* larvae was used as a template for probe synthesis. Fragments were amplified by PCR using the primer sets reported in Supplemental Table S5. The T3 promoter sequence and an EcoRI target site were added to the 5’ end of the forward primer, while the T7 promoter sequence and a KpnI target site were added to the 5’ end of the reverse primer. Amplicons were purified using the MinElute PCR Purification kit (Qiagen). Then, we digested the amplicons using EcoRI (antisense probe) or KpnI (sense probe), and added calf intestinal alkaline phosphatase (Promega) to prevent religation. DIG-labeled probes were prepared using a DIG RNA labeling mixture (Roche) and the T7 (antisense probe) or T3 RNA polymerase (sense probe). Synthesized probes were purified using the RNA Clean and Concentrator-5 kit (Zymo Research). The size of the probes was confirmed running an agarose gel.

### Fluorescence *in situ* hybridization

The FISH protocol used in this work is a slightly modified version of the protocol described in Jandura and colleagues (Jandura et al. 2017). Freshly dissected wing imaginal discs were fixed (30 min with 4% formaldehyde and 0.1% picric acid) and washed with PBTT (PBS with 0.1% Tween-20, 0.3% Triton X-100 and 0.1% picric acid). To quench endogenous peroxidase activity, the samples were incubated twice for 15 min with 0.3% hydrogen peroxidase in PBS and washed with PBTT. Then, samples were incubated with 80% acetone in PBS for 10 min at −20°C, washed with PBTT and rinsed with 1:1 PBTT:hybridization solution (50% formamide, 5X SSC buffer, 0.1% Tween-20, 0.3% Triton-X-100, 0.1 mg/ml of heparin and 1% salmon sperm ssRNA in DEPC water). Then, the samples were incubated with tempered pre-boiled hybridization solution for 3 hours at 56°C. The sense or antisense probe was diluted in hybridization solution (50 ng of probe per 100 µl of hybridization solution), and the mixture was denatured for 5 min at 85°C prior to the overnight incubation at 56°C. Then, probes were removed and the samples were washed for 15 min at 56°C with a decreasing concentration of hybridization solution:PBTT (3:1 twice, 1:1 and 1:3), followed by washes with PBTT. The samples were incubated with a blocking solution (1% BSA in PBTT) for 20 min at room temperature prior to the incubation with anti-digoxigenin (1:2,000 in blocking solution) for 2 hours. The anti-digoxigenin antibody was washed with blocking solution. For signal amplification purposes, the samples were incubated with the Tyramide Signal Amplification (TSA) system (Perkin Elmer) 2 hours in the dark, and then washed thoroughly with PBTT. Finally, samples were incubated for 15 min with YO-PRO-1 (Invitrogen), mounted in SlowFade Antifade (Life Technologies) and imaged under a fluorescence microscope.

### Image processing and analysis

Non-fluorescence images were taken using a Leica DMLB optical microscope, while fluorescence images were taken using a Leica SPE confocal microscope. All images were processed using Fiji (Schindelin et al. 2012) and the Adobe Illustrator software.

### Statistics

Prior to any statistical analysis, the Shapiro-Wilk normality test was used to check the data distribution. According to the result, the subsequent tests were parametric or non-parametric.

For comparisons of 2 groups, Student’s t-test or the Mann-Whitney U test were used. For comparisons of > 2 groups, one-way ANOVA or the Krustal-Wallis test followed by Dunn’s multiple comparisons test was used.

To address differences in the proportion of regenerated wings, a contingency table of regenerated and non-regenerated wings followed by the Fisher exact test was used. Bonferroni correction was applied when multiple comparisons were used.

All statistical tests were two-tailed. Differences were considered significant when *p*-values were less than 0.05 (*), 0.01 (**), or 0.001 (***). Tests were performed using GraphPad Prism 8 or R.

## Data access

All raw sequencing data generated in this study have been submitted to the NCBI Gene Expression Omnibus (GEO; https://www.ncbi.nlm.nih.gov/geo/) under accession number GSE223411.

## Acknowledgments

We thank Palmira Llorens-Giralt, Paula Climent-Cantó, José Esteban-Collado, Marta Morey, Elena Vizcaya-Molina, Marina Ruiz-Romero and Sílvia Pérez-Lluch for their insightful comments and suggestions, and Ignacio Maeso for his help on sequence conservation analysis. We thank the Bloomington *Drosophila* Stock Center (USA) for the fly stocks, FlyORF for the vector injection service, the Confocal Unit of the CCiT-UB for their technical help, and the Sequencing Unit of the CRG.

This project was funded by the following grants: PGC2018-099763-B100 and PID 2021-123300NB-I00 from the Spanish Government (Ministerio de Ciencia e Innovación); grant 2021SGR00293 from AGAUR (Generalitat de Catalunya); grant from the Institució Catalana de Recerca i Estudis Avançats (via an ICREA Academia award) to M.C.; New INDIGO grant to A.A.T. and M.C.. C.C.R. hold a predoctoral FPI contract (BES-2016-076691) from the Spanish Government (Ministerio de Ciencia e Innovación). R.A. hold a predoctoral fellowship of CONACYT, the “Becas al Extranjero” Program of Mexico.

## Author contributions

C.C.R. and M.C. conceived and designed the experiments. C.C.R. performed the experiments. M.T. and A.A.T. generated the mutants. C.C.R. and R.A.R. analyzed the data. M.C. supervised the project. M.C., F.S. and R.G. acquired the funding. C.C.R. and M.C. wrote the original draft. All authors reviewed the manuscript.

