## Supplemental Table S2 for "Expression dynamics of long non-coding RNAs during imaginal disc development and regeneration in *Drosophila*"

**Supplemental Table S2.** List of 131 differentially-expressed lncRNAs in regeneration.

| ncRNA | Type | Early | Mid | Late | Dev cluster |
|---|---|---|---|---|---|
| *CR31514* | Exonic | Upregulated | Non-DE | Non-DE | Cluster 4 |
| *CR32027* | Intronic | Non-DE | Non-DE | Upregulated | Cluster 1 |
| *CR32205* | Intergenic | Non-DE | Non-DE | Upregulated | Cluster 1 |
| *CR32207* | Intergenic | Upregulated | Upregulated | Upregulated | Cluster 1 |
| *CR32661* | Intergenic | Non-DE | Non-DE | Downregulated | Cluster 2 |
| *CR33940* | Exonic | Upregulated | Upregulated | Upregulated | Cluster 1 |
| *CR34046* | Intergenic | Downregulated | Non-DE | Non-DE | Cluster 3 |
| *CR34335* | Intronic | Downregulated | Downregulated | Downregulated | Cluster 4 |
| *CR40469* | Intergenic | Upregulated | Non-DE | Upregulated | Cluster 1 |
| *CR41257* | Intergenic | Upregulated | Non-DE | Upregulated | Cluster 2 |
| *CR42549* | Exonic | Non-DE | Non-DE | Downregulated | Cluster 2 |
| *CR42862* | Exonic | Non-DE | Downregulated | Non-DE | Cluster 4 |
| *CR42868* | Intergenic | Non-DE | Non-DE | Upregulated | Cluster 3 |
| *CR43144* | Intergenic | Downregulated | Downregulated | Downregulated | Cluster 3 |
| *CR43157* | Intergenic | Non-DE | Downregulated | Non-DE | Cluster 2 |
| *CR43264* | Intronic | Downregulated | Non-DE | Non-DE | Cluster 2 |
| *CR43436* | Intergenic | Non-DE | Downregulated | Downregulated | Cluster 3 |
| *CR43461* | Intronic | Downregulated | Downregulated | Downregulated | Cluster 2 |
| *CR43496* | Intronic | Downregulated | Downregulated | Downregulated | Cluster 2 |
| *CR43611* | Intronic | Upregulated | Upregulated | Upregulated | Cluster 3 |
| *CR43644* | Intergenic | Non-DE | Non-DE | Upregulated | Cluster 2 |
| *CR43684* | Intronic | Upregulated | Upregulated | Non-DE | Not expressed |
| *CR43837* | Intergenic | Downregulated | Non-DE | Non-DE | Cluster 3 |
| *CR43887* | Intergenic | Downregulated | Non-DE | Non-DE | Cluster 2 |
| *CR43900* | Exonic | Upregulated | Non-DE | Downregulated | Cluster 2 |
| *CR43918* | Intronic | Non-DE | Downregulated | Non-DE | Cluster 2 |
| *CR43927* | Intronic | Non-DE | Downregulated | Non-DE | Cluster 3 |
| *CR43943* | Intergenic | Non-DE | Downregulated | Non-DE | Cluster 2 |
| *CR43949* | Intergenic | Upregulated | Non-DE | Non-DE | Cluster 3 |
| *CR43956* | Intronic | Non-DE | Downregulated | Non-DE | Cluster 3 |
| *CR43963* | Exonic | Downregulated | Non-DE | Non-DE | Cluster 2 |
| *CR43971* | Exonic | Non-DE | Downregulated | Non-DE | Cluster 2 |
| *CR43995* | Intergenic | Downregulated | Downregulated | Downregulated | Cluster 2 |
| *CR44024* | Intergenic | Upregulated | Downregulated | Non-DE | Cluster 4 |
| *CR44029* | Exonic | Non-DE | Downregulated | Downregulated | Cluster 3 |
| *CR44042* | Intergenic | Non-DE | Non-DE | Downregulated | Cluster 2 |
| *CR44043* | Intronic | Non-DE | Upregulated | Non-DE | Cluster 3 |
| *CR44078* | Intergenic | Non-DE | Upregulated | Non-DE | Cluster 2 |
| *CR44119* | Exonic | Upregulated | Non-DE | Non-DE | Cluster 2 |
| *CR44127* | Intergenic | Downregulated | Downregulated | Non-DE | Cluster 2 |
| *CR44135* | Intergenic | Upregulated | Upregulated | Non-DE | Cluster 2 |
| *CR44172* | Intergenic | Downregulated | Non-DE | Non-DE | Cluster 2 |
| *CR44209* | Intergenic | Downregulated | Non-DE | Non-DE | Cluster 4 |

| | | | | | |
|---|---|---|---|---|---|
| *CR44294* | Intergenic | Upregulated | Non-DE | Non-DE | Cluster 3 |
| *CR44299* | Intronic | Non-DE | Non-DE | Downregulated | Cluster 2 |
| *CR44417* | Intronic | Non-DE | Downregulated | Non-DE | Cluster 3 |
| *CR44440* | Intergenic | Non-DE | Downregulated | Downregulated | Cluster 4 |
| *CR44458* | Intergenic | Non-DE | Downregulated | Non-DE | Cluster 4 |
| *CR44489* | Intergenic | Non-DE | Downregulated | Downregulated | Cluster 1 |
| *CR44499* | Intergenic | Downregulated | Non-DE | Non-DE | Cluster 1 |
| *CR44552* | Intergenic | Non-DE | Non-DE | Downregulated | Cluster 1 |
| *CR44589* | Intergenic | Non-DE | Downregulated | Non-DE | Cluster 4 |
| *CR44677* | Intergenic | Non-DE | Downregulated | Non-DE | Cluster 2 |
| *CR44749* | Intronic | Downregulated | Non-DE | Non-DE | Cluster 2 |
| *CR44761* | Exonic | Upregulated | Non-DE | Non-DE | Cluster 4 |
| *CR44784* | Intergenic | Non-DE | Non-DE | Upregulated | Cluster 2 |
| *CR44811* | Intronic | Non-DE | Non-DE | Downregulated | Cluster 3 |
| *CR44831* | Exonic | Non-DE | Downregulated | Non-DE | Cluster 2 |
| *CR44899* | Intergenic | Upregulated | Non-DE | Non-DE | Cluster 2 |
| *CR44917* | Intergenic | Non-DE | Non-DE | Upregulated | Cluster 3 |
| *CR44918* | Intergenic | Upregulated | Non-DE | Non-DE | Cluster 1 |
| *CR44955* | Intergenic | Upregulated | Upregulated | Non-DE | Cluster 2 |
| *CR44964* | Exonic | Non-DE | Upregulated | Non-DE | Cluster 2 |
| *CR44970* | Intergenic | Downregulated | Non-DE | Non-DE | Cluster 2 |
| *CR44971* | Intergenic | Downregulated | Non-DE | Non-DE | Cluster 1 |
| *CR44974* | Exonic | Upregulated | Non-DE | Non-DE | Cluster 2 |
| *CR44987* | Exonic | Non-DE | Downregulated | Downregulated | Cluster 3 |
| *CR44993* | Intronic | Upregulated | Downregulated | Non-DE | Cluster 3 |
| *CR45000* | Exonic | Non-DE | Non-DE | Upregulated | Cluster 2 |
| *CR45009* | Exonic | Downregulated | Non-DE | Non-DE | Cluster 2 |
| *CR45028* | Exonic | Upregulated | Non-DE | Non-DE | Cluster 2 |
| *CR45036* | Exonic | Non-DE | Downregulated | Non-DE | Cluster 2 |
| *CR45102* | Intergenic | Non-DE | Downregulated | Downregulated | Cluster 2 |
| *CR45128* | Intronic | Downregulated | Non-DE | Non-DE | Cluster 4 |
| *CR45171* | Exonic | Upregulated | Downregulated | Downregulated | Cluster 2 |
| *CR45176* | Exonic | Upregulated | Non-DE | Non-DE | Cluster 2 |
| *CR45181* | Intergenic | Non-DE | Downregulated | Non-DE | Cluster 2 |
| *CR45182* | Exonic | Downregulated | Non-DE | Non-DE | Cluster 3 |
| *CR45187* | Intergenic | Downregulated | Downregulated | Non-DE | Cluster 2 |
| *CR45232* | Intergenic | Non-DE | Downregulated | Non-DE | Cluster 1 |
| *CR45310* | Intergenic | Non-DE | Downregulated | Downregulated | Cluster 1 |
| *CR45311* | Intergenic | Non-DE | Downregulated | Non-DE | Cluster 1 |
| *CR45330* | Intergenic | Downregulated | Downregulated | Downregulated | Cluster 4 |
| *CR45433* | Intergenic | Upregulated | Non-DE | Non-DE | Cluster 2 |
| *CR45466* | Exonic | Non-DE | Downregulated | Non-DE | Cluster 2 |
| *CR45473* | Intergenic | Non-DE | Downregulated | Downregulated | Cluster 3 |
| *CR45479* | Exonic | Upregulated | Non-DE | Non-DE | Cluster 2 |
| *CR45485* | Exonic | Non-DE | Downregulated | Non-DE | Cluster 2 |
| *CR45501* | Exonic | Non-DE | Non-DE | Upregulated | Cluster 2 |

| | | | | | |
|---|---|---|---|---|---|
| *CR45517* | Intergenic | Non-DE | Downregulated | Non-DE | Cluster 4 |
| *CR45530* | Intergenic | Upregulated | Non-DE | Non-DE | Cluster 1 |
| *CR45550* | Intergenic | Non-DE | Downregulated | Non-DE | Cluster 1 |
| *CR45566* | Exonic | Non-DE | Non-DE | Upregulated | Cluster 1 |
| *CR45600* | Exonic | Non-DE | Downregulated | Downregulated | Cluster 2 |
| *CR45627* | Intergenic | Downregulated | Non-DE | Non-DE | Cluster 4 |
| *CR45638* | Intergenic | Downregulated | Non-DE | Non-DE | Cluster 1 |
| *CR45640* | Intergenic | Non-DE | Non-DE | Upregulated | Cluster 3 |
| *CR45641* | Intergenic | Downregulated | Non-DE | Non-DE | Cluster 2 |
| *CR45721* | Intergenic | Non-DE | Downregulated | Non-DE | Cluster 2 |
| *CR45734* | Intergenic | Downregulated | Downregulated | Non-DE | Cluster 1 |
| *CR45822* | Exonic | Non-DE | Downregulated | Downregulated | Cluster 2 |
| *CR45895* | Exonic | Non-DE | Downregulated | Non-DE | Cluster 2 |
| *CR45908* | Exonic | Non-DE | Downregulated | Non-DE | Cluster 4 |
| *CR45924* | Exonic | Downregulated | Non-DE | Non-DE | Cluster 4 |
| *CR45925* | Exonic | Non-DE | Downregulated | Upregulated | Cluster 4 |
| *CR46006* | Intergenic | Non-DE | Downregulated | Non-DE | Cluster 3 |
| *CR46044* | Exonic | Non-DE | Downregulated | Non-DE | Cluster 2 |
| *CR46056* | Intronic | Non-DE | Downregulated | Non-DE | Cluster 2 |
| *CR46057* | Intronic | Non-DE | Non-DE | Upregulated | Cluster 2 |
| *CR46093* | Exonic | Upregulated | Non-DE | Non-DE | Cluster 2 |
| *CR46123* | Intronic | Upregulated | Upregulated | Upregulated | Not expressed |
| *CR46139* | Exonic | Non-DE | Non-DE | Downregulated | Cluster 4 |
| *CR46194* | Exonic | Upregulated | Non-DE | Upregulated | Cluster 3 |
| *CR46252* | Intergenic | Downregulated | Non-DE | Non-DE | Cluster 2 |
| *CR46352* | Intergenic | Upregulated | Non-DE | Non-DE | Cluster 3 |
| *CR46410* | Intronic | Non-DE | Upregulated | Non-DE | Cluster 3 |
| *CR9284* | Intergenic | Non-DE | Non-DE | Downregulated | Cluster 2 |
| *dntRL* | Intergenic | Non-DE | Non-DE | Downregulated | Cluster 1 |
| *Hsromega* | Intergenic | Non-DE | Downregulated | Non-DE | Cluster 3 |
| *let7C* | Exonic | Non-DE | Downregulated | Non-DE | Cluster 2 |
| *noe* | Intronic | Non-DE | Downregulated | Non-DE | Cluster 4 |
| *RNaseP:RNA* | Intronic | Non-DE | Non-DE | Upregulated | Cluster 4 |
| *roX1* | Exonic | Non-DE | Non-DE | Downregulated | Cluster 2 |
| *roX2* | Intergenic | Downregulated | Downregulated | Downregulated | Cluster 4 |
| *sisRNA:1* | Intronic | Downregulated | Downregulated | Upregulated | Cluster 2 |
| *sisRNA:2* | Intronic | Downregulated | Non-DE | Non-DE | Cluster 4 |
| *sisRNA:3* | Intronic | Downregulated | Downregulated | Non-DE | Cluster 4 |
| *sisRNA:4* | Intronic | Non-DE | Downregulated | Non-DE | Cluster 2 |
| *sisRNA:CR46364* | Intronic | Downregulated | Non-DE | Non-DE | Cluster 2 |
| *Uhg3* | Intergenic | Non-DE | Non-DE | Downregulated | Cluster 2 |
| *Uhg8* | Exonic | Non-DE | Non-DE | Upregulated | Cluster 2 |
