## Supplemental Table S3 for "Expression dynamics of long non-coding RNAs during imaginal disc development and regeneration in *Drosophila*"

**Supplemental Table S3**. List of 131 DE lncRNA-PCG pairs and their classificatio

| ncRNA | ncRNA class | PCG | PCG class |
|---|---|---|---|
| *CR31514* | Valley | *CG12947* | Valley |
| *CR32027* | Peak | *Indy* | Peak |
| *CR32205* | Valley | *CG32206* | Not expressed |
| *CR32207* | Increasing | *CG32213* | Not expressed |
| *CR32661* | Valley | *pot* | Not expressed |
| *CR33940* | Increasing | *CG32212* | Valley |
| *CR34046* | Decreasing | *CG42693* | Valley |
| *CR34335* | Increasing | *DIP-alpha* | Not expressed |
| *CR40469* | Valley | *CG17636* | Not expressed |
| *CR41257* | Increasing | *rl* | Peak |
| *CR42549* | Peak | *mura* | Peak |
| *CR42862* | Decreasing | *CG33229* | Decreasing |
| *CR42868* | Valley | *CG42867* | Not expressed |
| *CR43144* | Decreasing | *Cul2* | Peak |
| *CR43157* | Decreasing | *Sclp* | Decreasing |
| *CR43264* | Peak | *CG15765* | Peak |
| *CR43436* | Decreasing | *betaGlu* | Peak |
| *CR43461* | Peak | *CG3168* | Peak |
| *CR43496* | Valley | *sov* | Increasing |
| *CR43611* | Peak | *CG45092* | Peak |
| *CR43644* | Valley | *E(spl)m3-HLH* | Decreasing |
| *CR43684* | Peak | *Sytbeta* | Not expressed |
| *CR43837* | Decreasing | *Inx2* | Not expressed |
| *CR43887* | Valley | *Dph1* | Not expressed |
| *CR43900* | Decreasing | *Cam* | Peak |
| *CR43918* | Decreasing | *Dp* | Peak |
| *CR43927* | Decreasing | *CG40006* | Peak |
| *CR43943* | Valley | *Hpd* | Decreasing |
| *CR43949* | Decreasing | *Notum* | Peak |
| *CR43956* | Decreasing | *anne* | Decreasing |
| *CR43963* | Decreasing | *Smr* | Not expressed |
| *CR43971* | Peak | *CG45071* | Increasing |
| *CR43995* | Increasing | *CG7133* | Peak |
| *CR44024* | Decreasing | *lgs* | Not expressed |
| *CR44029* | Decreasing | *unc-13* | Peak |
| *CR44042* | Decreasing | *l(2)41Ab* | Not expressed |
| *CR44043* | Peak | *l(2)41Ab* | Not expressed |
| *CR44078* | Peak | *TTLL4B* | Peak |
| *CR44119* | Valley | *SLC5A11* | Peak |
| *CR44127* | Increasing | *CG7881* | Not expressed |
| *CR44135* | Decreasing | *CG18547* | Peak |
| *CR44172* | Peak | *Tsp42Ef* | Peak |
| *CR44209* | Increasing | *Cyt-b5* | Not expressed |

| | | | |
|---|---|---|---|
| *CR44294* | Decreasing | *CG12128* | Peak |
| *CR44299* | Decreasing | *CG33144* | Peak |
| *CR44417* | Valley | *Mapmodulin* | Peak |
| *CR44440* | Peak | *PlexB* | Not expressed |
| *CR44458* | Valley | *Pms2* | Peak |
| *CR44489* | Valley | *CG43354* | Not expressed |
| *CR44499* | Valley | *mab-21* | Peak |
| *CR44552* | Decreasing | *mirr* | Peak |
| *CR44589* | Valley | *Plzf* | Peak |
| *CR44677* | Valley | *CG9300* | Peak |
| *CR44749* | Peak | *HmgD* | Peak |
| *CR44761* | Valley | *RYBP* | Peak |
| *CR44784* | Increasing | *CG30414* | Valley |
| *CR44811* | Peak | *bs* | Peak |
| *CR44831* | Peak | *pdgy* | Not expressed |
| *CR44899* | Decreasing | *CG13258* | Valley |
| *CR44917* | Valley | *tsh* | Peak |
| *CR44918* | Decreasing | *tsh* | Peak |
| *CR44955* | Decreasing | *Cyp12e1* | Increasing |
| *CR44964* | Peak | *CG14439* | Not expressed |
| *CR44970* | Peak | *kek1* | Peak |
| *CR44971* | Increasing | *kek1* | Peak |
| *CR44974* | Decreasing | *CycE* | Peak |
| *CR44987* | Peak | *galectin* | Peak |
| *CR44993* | Decreasing | *Df31* | Peak |
| *CR45000* | Valley | *Lcch3* | Not expressed |
| *CR45009* | Decreasing | *FucTC* | Not expressed |
| *CR45028* | Decreasing | *CG7702* | Increasing |
| *CR45036* | Peak | *CG6163* | Peak |
| *CR45102* | Peak | *Mnt* | Peak |
| *CR45128* | Valley | *Adf1* | Not expressed |
| *CR45171* | Decreasing | *Cpr67Fb* | Decreasing |
| *CR45176* | Decreasing | *Hpd* | Decreasing |
| *CR45181* | Valley | *vtd* | Peak |
| *CR45182* | Peak | *l(3)80Fg* | Valley |
| *CR45187* | Decreasing | *lost* | Peak |
| *CR45232* | Peak | *Atg9* | Increasing |
| *CR45310* | Decreasing | *CG46388* | Not expressed |
| *CR45311* | Valley | *CG46388* | Not expressed |
| *CR45330* | Decreasing | *CG5945* | Not expressed |
| *CR45433* | Decreasing | *Papss* | Peak |
| *CR45466* | Decreasing | *Cyp6v1* | Increasing |
| *CR45473* | Decreasing | *tyn* | Peak |
| *CR45479* | Decreasing | *Cyp4d14* | Valley |
| *CR45485* | Increasing | *CG3679* | Peak |
| *CR45501* | Valley | *parvin* | Peak |

| | | | |
|---|---|---|---|
| *CR45517* | Valley | *CG16756* | Not expressed |
| *CR45530* | Decreasing | *Ubi-p5E* | Peak |
| *CR45550* | Decreasing | *DNApol-iota* | Peak |
| *CR45566* | Valley | *RpL4* | Increasing |
| *CR45600* | Decreasing | *CG3984* | Not expressed |
| *CR45627* | Valley | *Pgant7* | Peak |
| *CR45638* | Increasing | *mtSSB* | Not expressed |
| *CR45640* | Increasing | *mtSSB* | Peak |
| *CR45641* | Decreasing | *mtSSB* | Peak |
| *CR45721* | Valley | *CG43354* | Not expressed |
| *CR45734* | Decreasing | *fng* | Peak |
| *CR45822* | Peak | *CTCF* | Peak |
| *CR45895* | Decreasing | *CG32163* | Decreasing |
| *CR45908* | Decreasing | *CG1943* | Not expressed |
| *CR45924* | Peak | *ana* | Not expressed |
| *CR45925* | Valley | *cer* | Decreasing |
| *CR46006* | Increasing | *CG12362* | Not expressed |
| *CR46044* | Valley | *Septin2* | Peak |
| *CR46056* | Decreasing | *E2f1* | Peak |
| *CR46057* | Valley | *E2f1* | Peak |
| *CR46093* | Decreasing | *BRWD3* | Peak |
| *CR46123* | Increasing | *Myo81F* | Not expressed |
| *CR46139* | Decreasing | *CG34112* | Decreasing |
| *CR46194* | Peak | *Oda* | Decreasing |
| *CR46252* | Decreasing | *vtd* | Peak |
| *CR46352* | Decreasing | *plh* | Not expressed |
| *CR46410* | Peak | *Ddr* | Peak |
| *CR9284* | Valley | *tpr* | Not expressed |
| *dntRL* | Decreasing | *CG17486* | Peak |
| *Hsromega* | Decreasing | *CG16791* | Peak |
| *let7C* | Peak | *CG10283* | Not expressed |
| *noe* | Valley | *blot* | Peak |
| *RNaseP:RNA* | Peak | *ATPsynC* | Peak |
| *roX1* | Decreasing | *ec* | Peak |
| *roX2* | Decreasing | *CG11695* | Increasing |
| *sisRNA:1* | Increasing | *Rga* | Increasing |
| *sisRNA:2* | Peak | *mbt* | Peak |
| *sisRNA:3* | Increasing | *Csp* | Peak |
| *sisRNA:4* | Increasing | *dpn* | Not expressed |
| *sisRNA:CR46364* | Decreasing | *RpS27* | Decreasing |
| *Uhg3* | Valley | *CG18810* | Not expressed |
| *Uhg8* | Valley | *atk* | Not expressed |

n.

| Pair class |
|---|
| Concordant |
| Concordant |
| Unrelated |
| Unrelated |
| Unrelated |
| Unrelated |
| Unrelated |
| Unrelated |
| Unrelated |
| Unrelated |
| Concordant |
| Concordant |
| Unrelated |
| Unrelated |
| Concordant |
| Concordant |
| Unrelated |
| Concordant |
| Unrelated |
| Concordant |
| Unrelated |
| Unrelated |
| Unrelated |
| Unrelated |
| Unrelated |
| Unrelated |
| Unrelated |
| Unrelated |
| Unrelated |
| Concordant |
| Unrelated |
| Unrelated |
| Unrelated |
| Unrelated |
| Unrelated |
| Unrelated |
| Unrelated |
| Concordant |
| Discordant |
| Unrelated |
| Unrelated |
| Concordant |
| Unrelated |

Unrelated
Unrelated
Discordant
Unrelated
Discordant
Unrelated
Discordant
Unrelated
Discordant
Discordant
Concordant
Discordant
Unrelated
Concordant
Unrelated
Unrelated
Discordant
Unrelated
Discordant
Unrelated
Concordant
Unrelated
Unrelated
Concordant
Unrelated
Unrelated
Unrelated
Discordant
Concordant
Concordant
Unrelated
Concordant
Concordant
Discordant
Discordant
Unrelated
Unrelated
Unrelated
Unrelated
Unrelated
Unrelated
Discordant
Unrelated
Unrelated
Unrelated
Discordant

| |
|---|
| Unrelated |
| Unrelated |
| Unrelated |
| Unrelated |
| Unrelated |
| Discordant |
| Unrelated |
| Unrelated |
| Unrelated |
| Unrelated |
| Unrelated |
| Concordant |
| Concordant |
| Unrelated |
| Unrelated |
| Unrelated |
| Unrelated |
| Discordant |
| Unrelated |
| Discordant |
| Unrelated |
| Unrelated |
| Concordant |
| Unrelated |
| Unrelated |
| Unrelated |
| Concordant |
| Unrelated |
| Unrelated |
| Unrelated |
| Unrelated |
| Discordant |
| Concordant |
| Unrelated |
| Discordant |
| Concordant |
| Concordant |
| Unrelated |
| Unrelated |
| Concordant |
| Unrelated |
| Unrelated |
