## Supplemental Table S4 for "Expression dynamics of long non-coding RNAs during imaginal disc development and regeneration in *Drosophila*"

**Supplemental Table S4**. List of differentially-expressed genes in *CR40469* mutants.

| Gene name | Gene type | Chromosome | Class |
|---|---|---|---|
| *CG5022* | mRNA | chr2L | Upregulated |
| *CG5096* | mRNA | chr2L | Upregulated |
| *CR44119* | nRNA | chr2L | Upregulated |
| *Fbp2* | mRNA | chr2L | Upregulated |
| *G6P* | mRNA | chr2L | Upregulated |
| *CG9877* | mRNA | chr2R | Upregulated |
| *CR44042* | nRNA | chr2R | Upregulated |
| *GstE8* | mRNA | chr2R | Upregulated |
| *Pu* | mRNA | chr2R | Upregulated |
| *SkpB* | mRNA | chr2R | Upregulated |
| *Spn47C* | mRNA | chr2R | Upregulated |
| *St3* | mRNA | chr2R | Upregulated |
| *Apl* | mRNA | chr3L | Upregulated |
| *Cpr66D* | mRNA | chr3L | Upregulated |
| *CR45181* | nRNA | chr3L | Upregulated |
| *NijA* | mRNA | chr3L | Upregulated |
| *tut* | mRNA | chr3L | Upregulated |
| *AstA* | mRNA | chr3R | Upregulated |
| *CG12224* | mRNA | chr3R | Upregulated |
| *CG31075* | mRNA | chr3R | Upregulated |
| *CG33099* | mRNA | chr3R | Upregulated |
| *CG6654* | mRNA | chr3R | Upregulated |
| *CR44291* | nRNA | chr3R | Upregulated |
| *CR45641* | nRNA | chr3R | Upregulated |
| *Orct* | mRNA | chr3R | Upregulated |
| *p24-2* | mRNA | chr3R | Upregulated |
| *pinta* | mRNA | chr3R | Upregulated |
| *tobi* | mRNA | chr3R | Upregulated |
| *Unc-115b* | mRNA | chr3R | Upregulated |
| *ABCA* | mRNA | chrX | Upregulated |
| *AgmNAT* | mRNA | chrX | Upregulated |
| *cac* | mRNA | chrX | Upregulated |
| *CG15370* | mRNA | chrX | Upregulated |
| *CG15892* | mRNA | chrX | Upregulated |
| *CG32631* | mRNA | chrX | Upregulated |
| *CG32640* | mRNA | chrX | Upregulated |
| *CG3603* | mRNA | chrX | Upregulated |
| *CG4313* | mRNA | chrX | Upregulated |
| *CG6023* | mRNA | chrX | Upregulated |
| *CG6999* | mRNA | chrX | Upregulated |
| *CR43496* | nRNA | chrX | Upregulated |
| *CR44964* | nRNA | chrX | Upregulated |
| *CR45466* | nRNA | chrX | Upregulated |

| | | | |
|---|---|---|---|
| *CR45485* | nRNA | chrX | Upregulated |
| *Gbeta5* | mRNA | chrX | Upregulated |
| *png* | mRNA | chrX | Upregulated |
| *ppk28* | mRNA | chrX | Upregulated |
| *t* | mRNA | chrX | Upregulated |
| *CG31809* | mRNA | chr2L | Downregulated |
| *Nepl3* | mRNA | chr2L | Downregulated |
| *pes* | mRNA | chr2L | Downregulated |
| *SoYb* | mRNA | chr2L | Downregulated |
| *CG11400* | mRNA | chr2R | Downregulated |
| *CG15236* | mRNA | chr2R | Downregulated |
| *CG43103* | mRNA | chr2R | Downregulated |
| *CG6701* | mRNA | chr2R | Downregulated |
| *Cpr47Ee* | mRNA | chr2R | Downregulated |
| *CR43907* | nRNA | chr2R | Downregulated |
| *Cyp6a17* | mRNA | chr2R | Downregulated |
| *Ir56b* | mRNA | chr2R | Downregulated |
| *vis* | mRNA | chr2R | Downregulated |
| *CG32110* | mRNA | chr3L | Downregulated |
| *CG32368* | mRNA | chr3L | Downregulated |
| *CG5282* | mRNA | chr3L | Downregulated |
| *CR32205* | nRNA | chr3L | Downregulated |
| *CR32207* | nRNA | chr3L | Downregulated |
| *CR33940* | nRNA | chr3L | Downregulated |
| *CR43426* | nRNA | chr3L | Downregulated |
| *CR45433* | nRNA | chr3L | Downregulated |
| *NT1* | mRNA | chr3L | Downregulated |
| *Act87E* | mRNA | chr3R | Downregulated |
| *CG31262* | mRNA | chr3R | Downregulated |
| *CG5217* | mRNA | chr3R | Downregulated |
| *CR44955* | nRNA | chr3R | Downregulated |
| *CR46107* | nRNA | chr3R | Downregulated |
| *RpS19b* | mRNA | chr3R | Downregulated |
| *unc-13* | mRNA | chr4 | Downregulated |
| *CG2898* | mRNA | chrX | Downregulated |
| *CG3168* | mRNA | chrX | Downregulated |
| *CG3568* | mRNA | chrX | Downregulated |
| *CG42259* | mRNA | chrX | Downregulated |
| *CG43672* | mRNA | chrX | Downregulated |
| *CG4607* | mRNA | chrX | Downregulated |
| *CR40469* | nRNA | chrX | Downregulated |
| *CR44957* | nRNA | chrX | Downregulated |
| *CR45479* | nRNA | chrX | Downregulated |
| *Crg-1* | mRNA | chrX | Downregulated |
| *Cubn* | mRNA | chrX | Downregulated |
| *Cyp4d14* | mRNA | chrX | Downregulated |

| | | | |
|---|---|---|---|
| *dpr18* | mRNA | chrX | Downregulated |
| *Karl* | mRNA | chrX | Downregulated |
| *Lgr4* | mRNA | chrX | Downregulated |
| *Nep1* | mRNA | chrX | Downregulated |
| *Pde9* | mRNA | chrX | Downregulated |
| *Sdic2* | mRNA | chrX | Downregulated |
