## Supplemental Table S5 for "Expression dynamics of long non-coding RNAs during imaginal disc development and regeneration in *Drosophila*"

**Supplemental Table S5**. List of oligonucleotides used in this \

| Region | Forward sequence (5'-->3') |
|---|---|
| Homology arm 1 (*CR40469* deletion) | GAGGAATTCGAACTGGACATCATGCGTTG |
| Homology arm 2 (*CR40469* deletion) | GAGACTAGTTGGCAACAACATCTTGAACC |
| Control primers for deletion validation | CATTGCATGAGACGATGGCG |
| Deletion primers for deletion validation | TTGTGCGCAGAGCATTGACTT |
| RNA probe synthesis | TAATACGACTCACTATAGGGGGAATTCCCACGTTCC TCACTAATTGTGGCTA |
| *CR40469* transgene amplification | CGAAGAATTCCACGTTCCTCACTAATTGTGGCTA |
| *CR34335* transgene amplification | CGAAGAATTCGGCGGTCGAGTGCCTCACA |
| *AstA-R1* | ATGTGGTGGCGTACACGAAT |
| *CG13021* | TCCCCAAGGAAACTCTACTGTG |
| *CG17636* | AGTGGGAGTGGAGCGGCATT |
| *CG17707* | TGCGGTTTTTACTTTCATCGG |
| *CG2875* | GCCAAACCGAAAATGAAACCCT |
| *CG45473* | GATGTTGGTGAACGCGGTACT |
| *CR34335* | TTGCATGAGACGATGGCGCA |
| *CR40469/CR34335* | TTGAGTTCCTCCAACACAGCGT |
| *CR43863* | CCACTTGGCGTGACGAAATC |
| *CR44999* | TGCAACGATATGGTGGACGA |
| *DIP-a* | TTTCGTTGCCATCTTCGCGGC |
| *Gad8* | CGGGAGGAGCGTAACTACTTC |
| *ppk8* | TCCATTGCCAACGAGCCTTT |
| *RhoGAP1A* | ATATTCGGAGGCCACCGCTC |
| *sply* | CTTTCCCGATTCCCGTA |
| *tyn* | ACCACCGGGAACTCTGGATA |

work.

| Reverse sequence (5'-->3') |
|---|
| GAGGCGGCCGCTGCTGGCATTATGTTGGTGT |
| GAGAGATCTTGTAACACGCACAAGGAAGC |
| TTTCCTCTCCCTACTGCACA |
| GGAAAATCTCCCAGCACACAAA |
| AATTAACCCTCACTAAAGGGGGGTACCCCGGAT TTTTTGCTTGTTTTGTTTGA |
| GGATTTTTTGCTTGTTTTGTTTGA |
| AAAAAAAAGCGATCAGACTGGAATGTGAGAA |
| TTGCAGTTCGCGTTGTCTTG |
| GCCAGCTGATCCGTCAAGTT |
| CAGGCAATGGGCGGCATTGTT |
| ACAGACCGGCTAAATCTGCAA |
| GCTGGCATATTATCGGGCGT |
| AGCTTTTGCTTATGAACGCGGG |
| CGGTCGAGTGCCTCACAGTGTA |
| TTGCGCCATCGTCTCATGCAA |
| CCGCGTATGTTTGCTATGCC |
| CACTTTGCCGTTACTCCTGC |
| AGGGCAGTTCCGTGCGATTGA |
| GCTGCATCACATGCTTGGTA |
| GTTCAACGTCCACCTTTCGC |
| CTGGCTGGGAATGGTAGGGG |
| TGACGGGCTTAAGGCAATC |
| CGGACCTGTACGGCAATTCT |
