## Supplemental Table S6 for "Expression dynamics of long non-coding RNAs during imaginal disc development and regeneration in *Drosophila*"

**Supplemental Table S6**. Summary of sequenced samples and mapped reads for the total RNA

| Sample name | Genotype | Origin | Time point | Library |
|---|---|---|---|---|
| Control rep1 | *sal*$^{E/Pv}$ *-LHG:tub-Gal80*$^{TS}$ *; lexO-rpr* | Wing disc | Early regeneration | total RNA-seq |
| Control rep2 | | Wing disc | Early regeneration | total RNA-seq |
| Control rep3 | | Wing disc | Early regeneration | total RNA-seq |
| Mutant rep1 | *CR40469*$^{-/-}$ *; sal*$^{E/Pv}$ *-LHG:tub-Gal80*$^{TS}$ *; lexO-rpr* | Wing disc | Early regeneration | total RNA-seq |
| Mutant rep2 | | Wing disc | Early regeneration | total RNA-seq |
| Mutant rep3 | | Wing disc | Early regeneration | total RNA-seq |

\-seq.

| Mapped reads |
| --- |
| 44522506 |
| 45405499 |
| 45899544 |
| 44361094 |
| 45919096 |
| 47365674 |
