## Supplemental Fig. S1 for "Expression dynamics of long non-coding RNAs during imaginal disc development and regeneration in *Drosophila*"

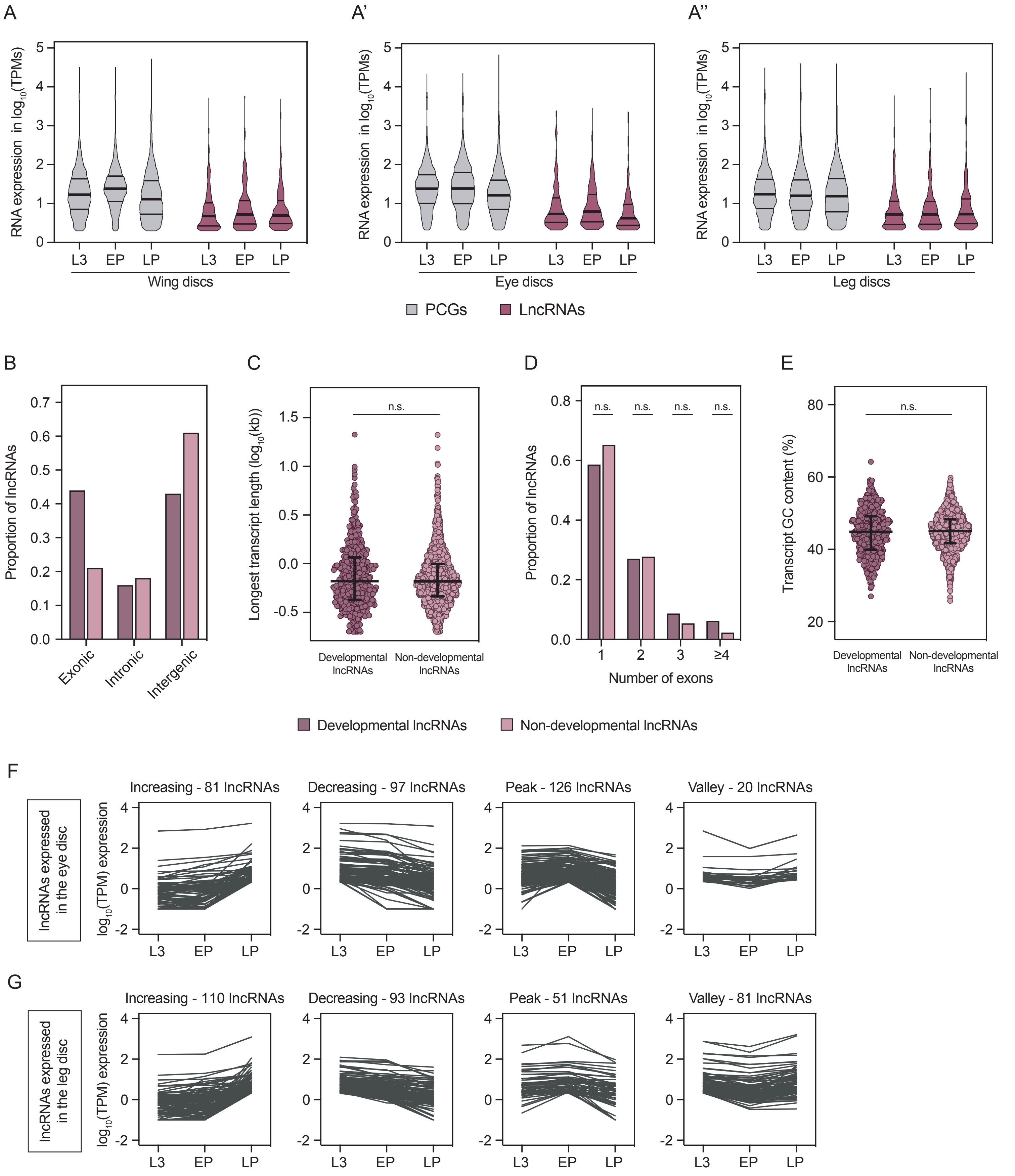


**Supplemental Fig. S1.** Characterization of genes expressed in developing imaginal discs. (**A-A'-A'''**) Violin plots showing the expression of PCGs and lncRNAs expressed at least 1 TPM in the wing, eye or leg discs. The first quartile, median and third quartile are represented. (**B**) Classification of developmental lncRNAs (expressed in at least one developmental sample) and non-developmental lncRNAs (not expressed in the wing, eye and leg discs) into exonic, intronic or intergenic. (**C**) Length of the longest transcript for each lncRNA. Lenght is presented as the log base 10 in kilobases. Median and interquartile range are represented. (**D**) Number of exons of the longest transcript for each lncRNA. (**E**) GC content of the longest transcript for each lncRNA. Median and interquartile range are represented. (**F**) Classification of the lncRNAs expressed in the eye disc during development. (**G**) Classification of the lncRNAs expressed in the leg disc during development. Data from Ruiz-Romero et al. 2022.
