## Supplemental Fig. S2 for "Expression dynamics of long non-coding RNAs during imaginal disc development and regeneration in *Drosophila*"

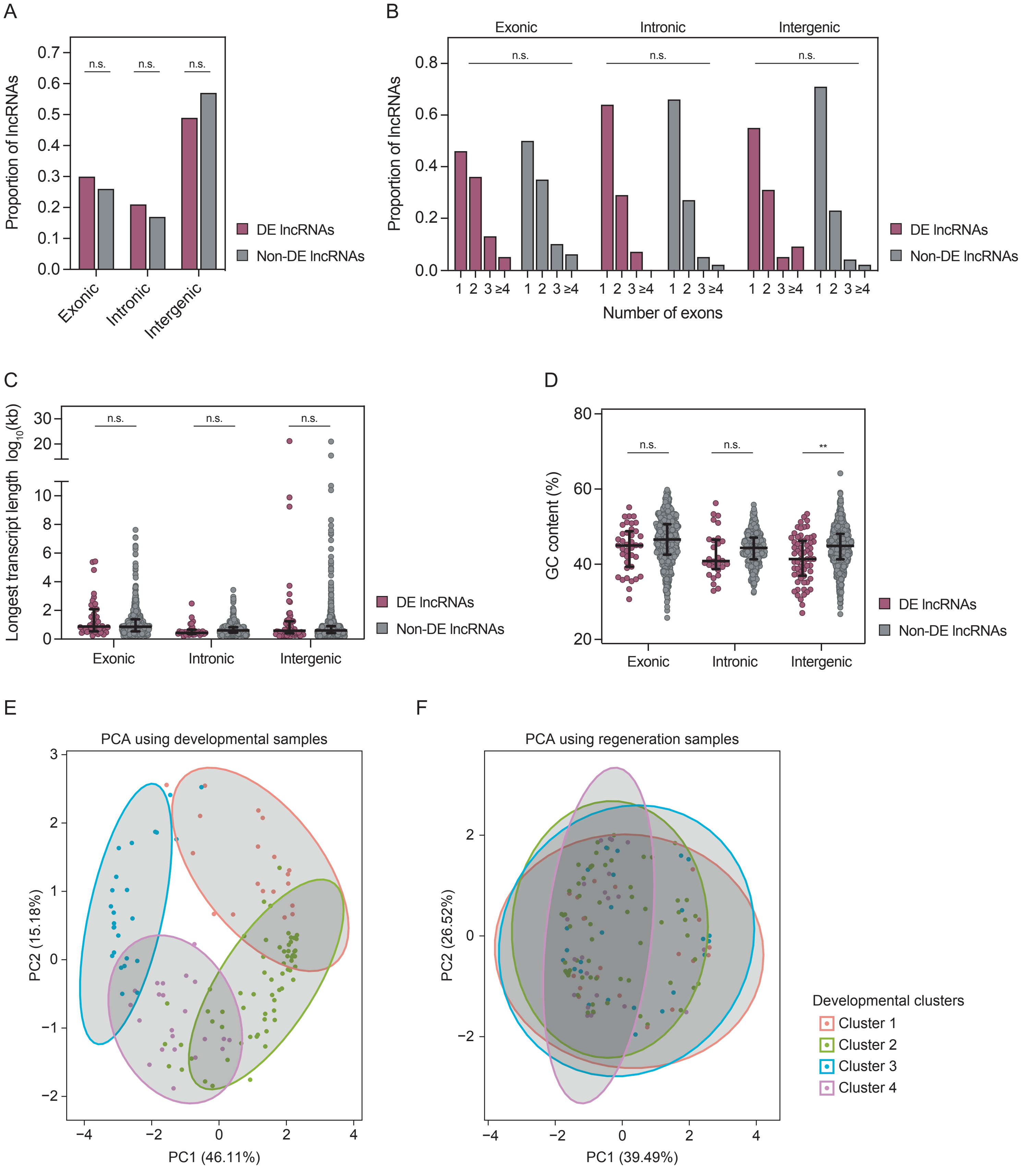


**Supplemental Fig. S2.** Differentially-expressed (DE) lncRNAs in regeneration. (**A**) Proportion of exonic, intronic and intergenic lncRNAs. (**B**) Classification of exonic, intronic and intergenic lncRNAs according to the number of exons in their longest transcript. (**C**) Longest transcript length of exonic, intronic and intergenic lncRNAs. (**D**) GC content of the longest transcript of exonic, intronic and intergenic lncRNAs. (**E**) Principal component analysis (PCA) based on the expression in development of DE lncRNAs. (**F**) PCA based on the expression in regeneration of DE lncRNAs. The color code in E and F represents the 4 gene clusters defined in Figure 3H. PCAs shown in E and F were based on the expression of the 129 clustered lncRNAs differentially-expressed in regeneration normalized as Z-scores. The median and interquartile range are represented in C and D. $p < 0.01$ (**); n.s. = non-significant. Data from Vizcaya-Molina et al 2018.
