## Supplemental Fig. S3 for "Expression dynamics of long non-coding RNAs during imaginal disc development and regeneration in *Drosophila*"

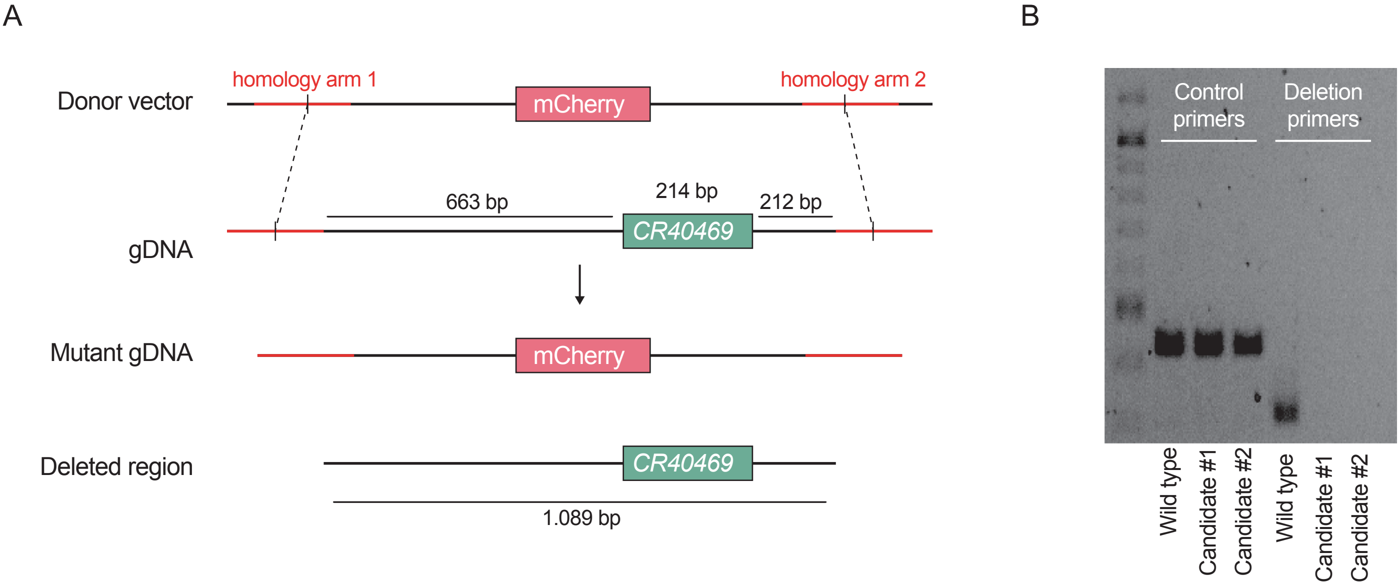


C

```
CR40469         -------------------------------------CACGTTCCTCACTAATTGTGGCTA  24
CR34335-short   GGCGGTCGAGTGCCTCACAGTGTATCAAGGGTTGGCCACGTTCCTCACTAATTGTGGCTA  60
CR34335-long    GGCGGTCGAGTGCCTCACAGTGTATCAAGGGTTGGCCACGTTCCTCACTAATTGTGGCTA  60
                                                     ************************

CR40469         TTTGCGCCATCGTCTCATGCAATGTTATTTGAGAGATGGCAAAATATATAGTATGTTTGT  84
CR34335-short   TTTGCGCCATCGTCTCATGCAATGTTATTTGAGAGATGGCAAAATATATAGTATGTTTGT  120
CR34335-long    TTTGCGCCATCGTCTCATGCAATGTTATTTGAGAGATGGCAAAATATATAGTATGTTTGT  120
                ************************************************************

CR40469         CTCCAATGTGTTGAGACTGAGAAGATATTGTACCCGTGAATTGATGAAAATTGATTGATT  144
CR34335-short   CTCCAATGTGTTGAGACTGAGAAGATATTGTACCCGTGAATTGATGAAAATTGATTGATT  180
CR34335-long    CTCCAATGTGTTGAGACTGAGAAGATATTGTACCCGTGAATTGATGAAAATTGATTGATT  180
                ************************************************************

CR40469         ATATTGTAATGTTGATTTCATGAAAAACACGCTGTGTTGGAGGAACTCAAACAAAACAAG  204
CR34335-short   ATAT-GTAATGTTGATTTCATGAAAAACACGCTGTGTTGGAGGAACTCAAACAAAACAAG  239
CR34335-long    ATAT-GTAATGTTGATTTCATGAAAAACACGCTGTGTTGGAGGAACTCAAACAAAACAAG  239
                **** *******************************************************

CR40469         CAAAAAATCC--------------------------------------------------  214
CR34335-short   CATAAAATCC--------------------------------------------------  249
CR34335-long    CATAAAATCCAAAAAAAAAAAAAAAAAAAACAAATCAAATTTTAACAAACAATAATAATAC  299
                ** *******

CR40469         ------------------------------------------------------------  214
CR34335-short   ------------------------------------------------------------  249
CR34335-long    TGTGTGGTGCGCTGGCGTGGGGGAGTGGTTATTCTCACATTCCAGTCTGATCGCTTTTTT  359


CR40469         --   214
CR34335-short   --   249
CR34335-long    TT   361


Identities 212/214(99%)
Gaps 1/214(0%)
```

**Supplemental Fig. S3.** *CR40469* mutant validation and sequence alignment. (**A**) Schematic representation of the deleted genomic region containing the full *CR40469* locus, 663 bp upstream and 212 bp downstream. (**B**) Electrophoresis of PCR products to confirm the deletion of *CR40469* . Primers hybridizing 3 Mb downstream of the deletion site were used as control primers, while primers hybridizing within the deleted region were used to confirm the absence of the deleted sequence. The deletion was confirmed for candidates #1 and #2. (**C**) Sequence alignment of the *CR40469* and *CR34335* transcripts. The single transcript of *CR40469* and the short and long isoforms of *CR34335* were used for the alignment. 212 of the 214 nt of the *CR40469* transcript are identical in both *CR34335* isoforms. Identities are marked with an asterisk (*).
