## Supplemental Fig. S4 for "Expression dynamics of long non-coding RNAs during imaginal disc development and regeneration in *Drosophila*"

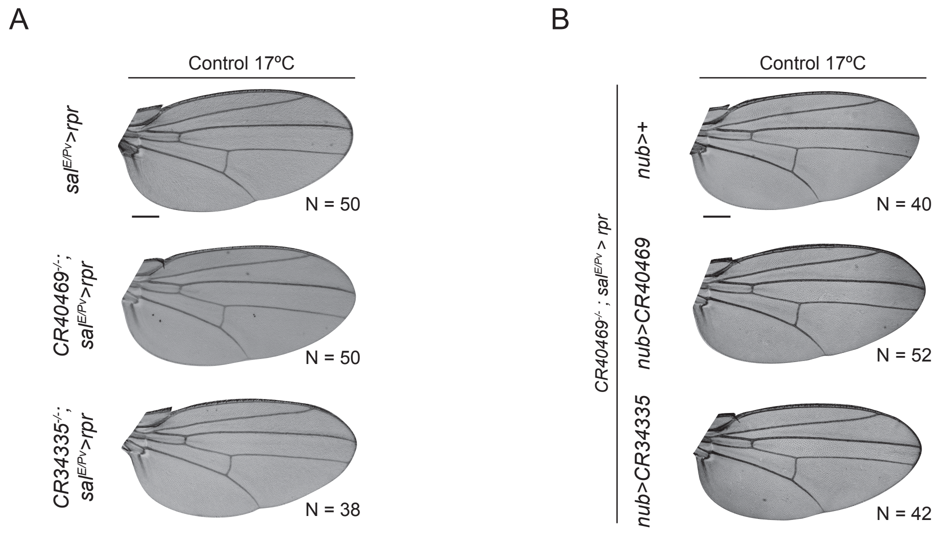


**Supplemental Fig. S4.** Sample images from adult wings incubated at 17ºC. (**A**) Control adult wings containing *sal$^{E/Pv}$>rpr* in a wild type, *CR40469* homozygous mutant, or *CR34335* homozygous mutant background. (**B**) Control adult wings containing *sal$^{E/Pv}$>rpr* in a *CR40469* homozygous mutant background, plus *nub*>+, *nub>CR40469* or *nub>CR34335.*
