## Supplemental Fig. S5 for "Expression dynamics of long non-coding RNAs during imaginal disc development and regeneration in *Drosophila*"

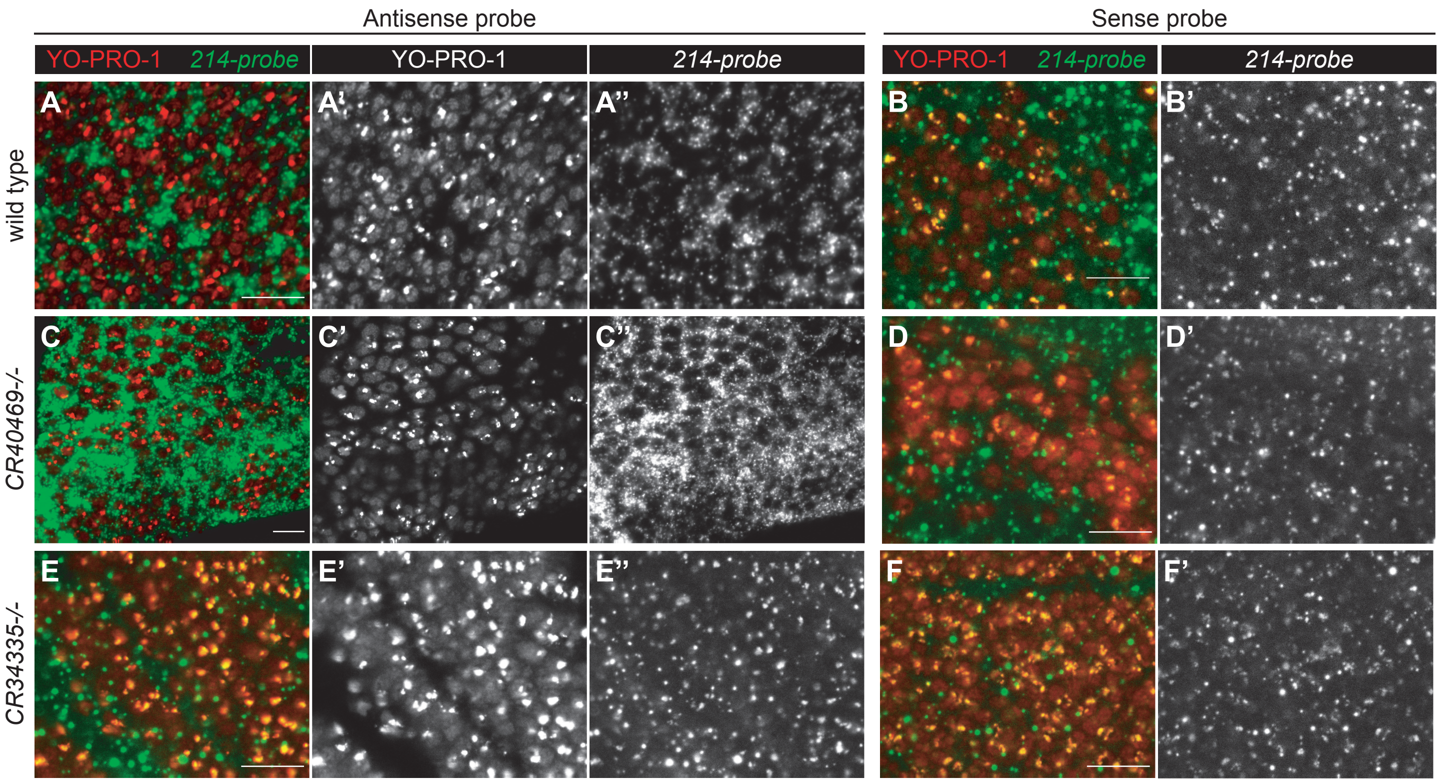


**Supplemental Fig. S5.** Subcellular localization of *CR40469* and *CR34335* in the wing disc. FISH imaging of (**A-B**) wild type, (**C-D**) *CR40469* homozygous mutants, and (**E-F**) *CR34335* homozygous mutants hybridized with the antisense *214-probe* designed to detect the *CR40469* and *CR34335* transcripts (A,C,E), or hybridized with a sense *214-probe* as negative control (B,D,F). N ≥ 10 per condition. Scale bar = 10 µm.
