## Supplemental Fig. S6 for "Expression dynamics of long non-coding RNAs during imaginal disc development and regeneration in *Drosophila*"

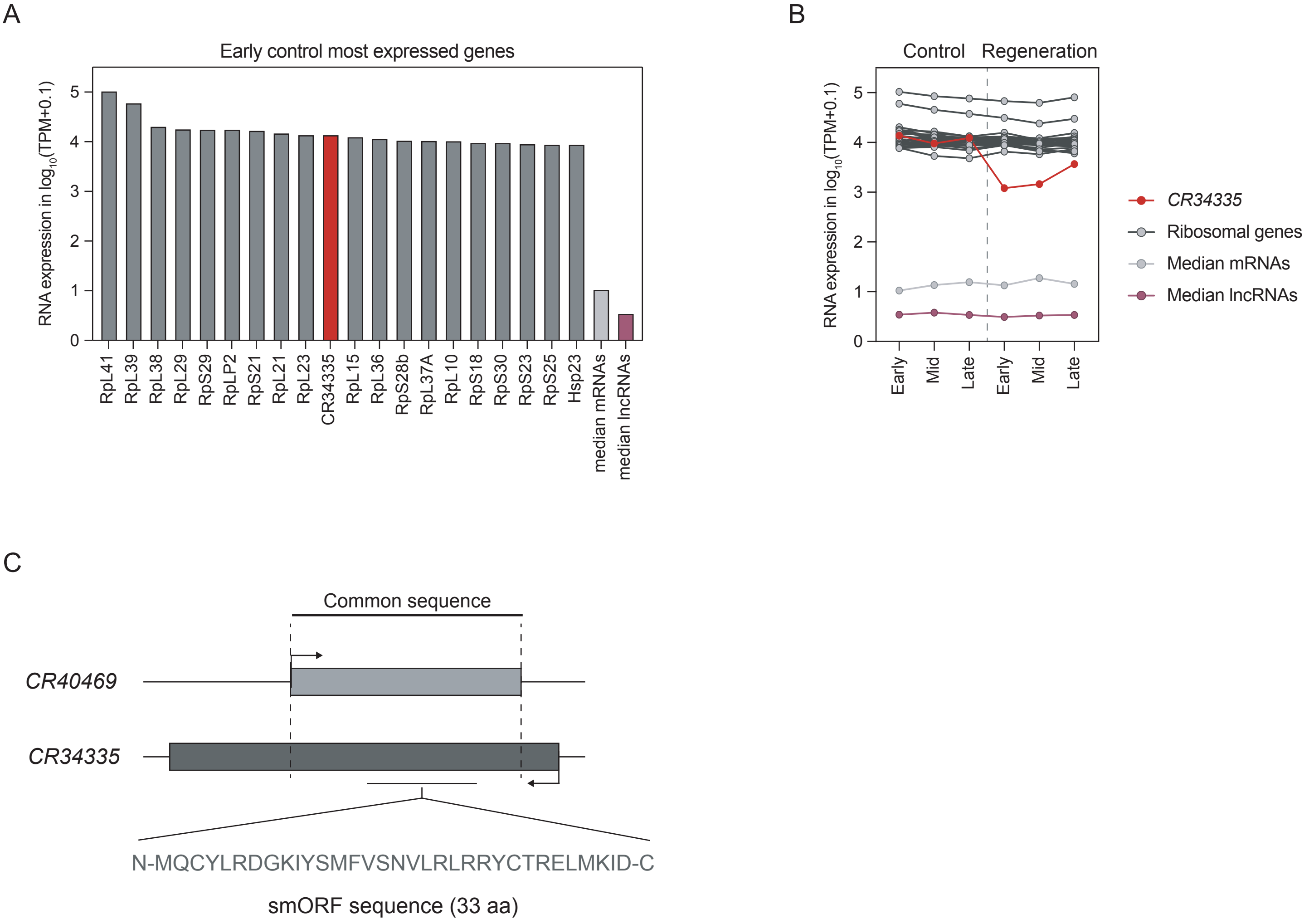


**Supplemental Fig. S6.** Analysis of *CR34335* expression and putative small ORF. (**A**) Expression of the 20 most expressed genes in the early control wing discs. (**B**) Expression of *CR34335* compared to that of ribosomal genes in the control and regeneration samples during the early, mid and late stages. (**C**) Predicted small ORF within the *CR40469* and *CR34335* RNA sequence using ORFinder. Localization and amino acid sequence of the predicted small ORF present in the *CR40469* and *CR34335* transcripts. Data from Vizcaya-Molina et al. 2018.
