## Supplemental Fig. S7 for "Expression dynamics of long non-coding RNAs during imaginal disc development and regeneration in *Drosophila*"

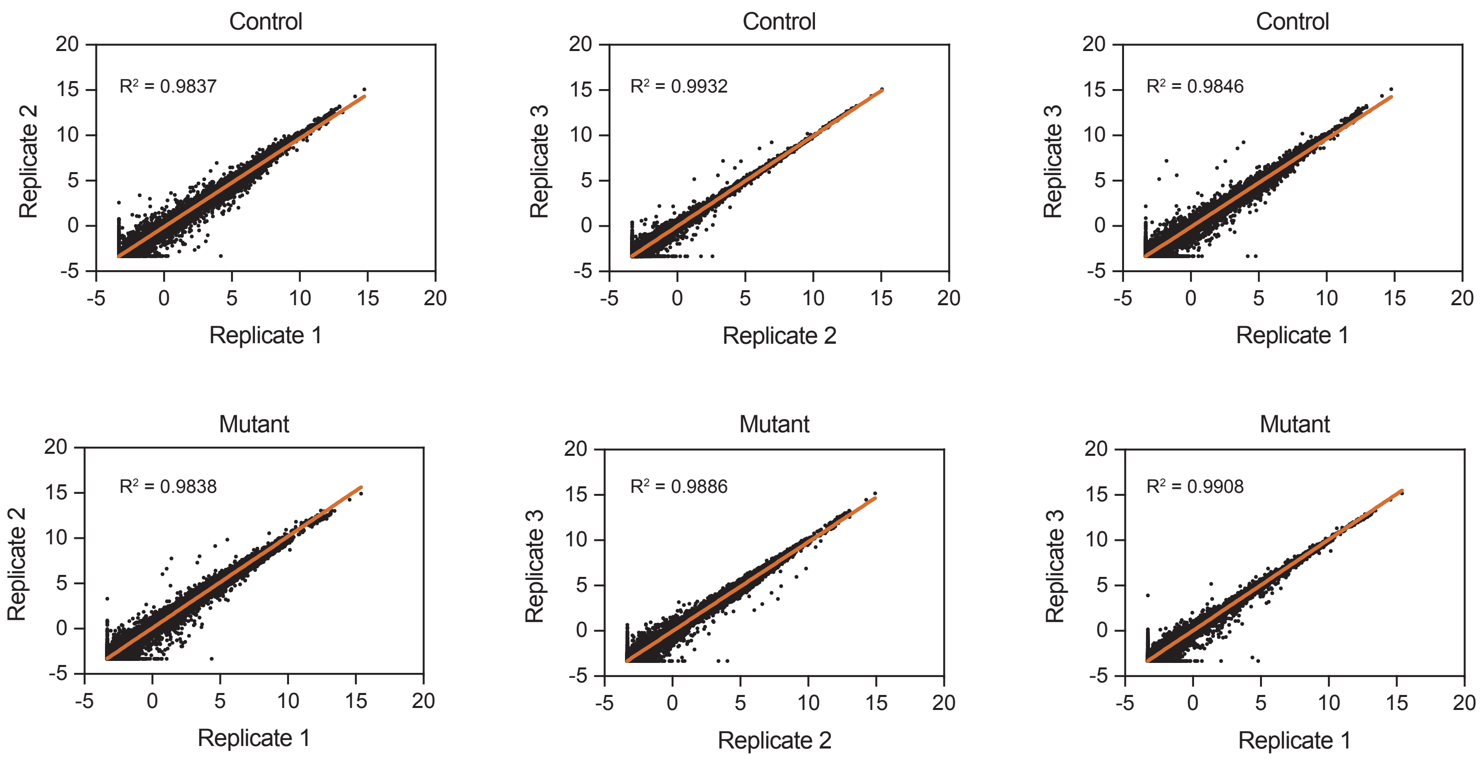


**Supplemental Fig. S7.** Replicate analysis of RNA-seq data. Scatter plots showing the correlation between the three biological replicates per each condition. Each dot represents the -$\log_2$ TPMs of each gene per replicate. Coefficient of determination ($R^2$) was higher than 0.98 in each comparison. The linear regression line is represented in orange.
